# Molecular competition in G1 controls when cells simultaneously commit to terminally differentiate and exit the cell-cycle

**DOI:** 10.1101/632570

**Authors:** Michael L. Zhao, Atefeh Rabiee, Kyle M. Kovary, Zahra Bahrami-Nejad, Brooks Taylor, Mary N. Teruel

## Abstract

Terminal differentiation is essential for the development and maintenance of tissues in all multi-cellular organisms and is associated with a permanent exit from the cell cycle. Failure to permanently exit the cell cycle can result in cancer and disease. However, the molecular mechanisms and timing that coordinates differentiation commitment and cell cycle exit are not yet understood. Here using adipogenesis as a model system to track differentiation commitment in live cells, we show that a rapid switch mechanism engages exclusively in G1 to trigger a simultaneous commitment to differentiate and permanently exit from the cell cycle. We identify a signal integration mechanism whereby the strengths of both mitogen and differentiation stimuli control a molecular competition between cyclin D1 and PPARG-induced expression of the CDK inhibitor p21 which in turn regulates if and when the differentiation switch is triggered and when the proliferative window closes. In this way, the differentiation control system is able to couple mitogen and differentiation stimuli to sustain a long-term balance between terminally differentiating cells and maintaining the progenitor cell pool, a parameter of critical importance for enabling proper development of tissue domains and organs.

**HIGHLIGHTS:**

- Progenitor cells both commit to terminally differentiate and permanently exit the cell cycle at a precise time in G1 as a result of a competition process that can last over multiple cell cycles.
- Positive-feedback driven expression of PPARG and the parallel induction of p21 triggers a rapid commitment to terminally differentiate and then maintains a postmitotic adipocyte state.
- Opposing mitogen and adipogenic signals are funneled into a molecular competition in G1 phase that controls if and when cells commit to differentiate, which in turn regulates the number of differentiated cells produced while allowing for the maintenance of sufficient progenitor cells.

---

Terminal differentiation is essential for developing, maintaining, and regenerating tissues and is the mechanism by which neurons, skeletal muscle cells, adipocytes (fat cells), and many other critical cell types are generated in humans and other multi-cellular organisms (Ruijtenberg and van den Heuvel, 2016). Terminal differentiation typically requires that proliferative progenitor cells transition into post-mitotic differentiated cells that remain in a permanent state of withdrawal from the cell cycle. Failure of terminally differentiated cells to enter and maintain the post-mitotic state can lead to disease and is a hallmark of cancer (Ghaben and Scherer, 2019; Ruijtenberg and van den Heuvel, 2016). However, despite the fundamental importance of coordinating cell-cycle exit and terminal differentiation for human health, when and how permanent cell cycle exit is achieved relative to terminal differentiation and whether terminal differentiation is a parallel but independent process that can occur before or after permament cell-cycle exit remained unclear (Buttitta and Edgar, 2007; Hardwick et al., 2015; Soufi and Dalton, 2016).

The question we are addressing in our study arises from preceding studies that found different molecular links between the cell cycle and terminal differentiation. For example, links between cyclin-dependent kinase (CDK) inhibitors and differentiation have been reported in terminally differentiated cells, suggesting parallel pathways as well as tissue-specific redundancy in coordinating cell cycle exit and terminal differentiation (Buttitta et al., 2007; Parker et al., 2006; Ruijtenberg et al., 2015; Zalc et al., 2014). Also, lengthening of the G1-phase of the cell cycle has been observed during differentiation of diverse cell types, including human embryonic stem cells (Calder et al., 2012), neuroendocrine cells (Krentz et al., 2017; Miyatsuka et al., 2011), and neurons (Lange et al., 2009). However, even though such molecular links have been found, there is conflicting evidence on whether and how strongly terminal differentiation and the cell cycle are connected. For example, studies in adipocytes and neurons suggested that G1-lengthening, cell cycle exit, and terminal differentiation happen sequentially (Lange et al., 2009; Tang et al., 2003) whereas other studies support a model that cell cycle exit and terminal differentiation occur as parallel, independently-regulated processes (Lacomme et al., 2012; Qiu et al., 2001). Still other studies suggest that the two processes are coordinated by dual actions of core components of the cell cycle and differentiation machinery (Hardwick et al., 2015).

One challenge in understanding the relationship between cell-cycle exit and terminal differentiation is that there is great variability in whether and when individual progenitor cells in the same population proliferate or differentiate during the several-day long differentiation process. This variation between cells makes it for example difficult to answer the simple question how many cell divisions occur before cells terminally differentiate. To overcome this challenge, methods are needed that can measure whether and when during the multi-day differentiation timecourse an individual cell commits to irreversibly differentiate. Such methods require being able to track both cell cycle and differentiation progression simultaneously in realtime. Live-cell imaging approaches have been used in stem cells that undergo a slowing or transient exit from the cell cycle during the differentiation process (Matson et al., 2017; Pauklin and Vallier, 2013). However, live-cell imaging studies to uncover the link between the cell cycle and terminal cell differentiation have to our knowledge not yet been made, and the underlying regulatory mechanisms are likely different when permanent exit from, rather than temporary slowing of the cell cycle is required.

Many terminal cell differentiation processes including adipogenesis and myogenesis are regulated by a cascade of transcription factors (Blais et al., 2005; Farmer, 2006). In order to establish a temporal marker for differentiation commitment, it is essential to determine which of the factors in the transcriptional cascade controlling the terminal differentiation process exhibits bimodal and irreversible behavior and can thus be used to distinguish whether a cell still has the option to remain undifferentiated or has committed to terminally differentiate. For example, in adipogenesis, even though the early transcription factor C/EBPB is required, it is not a suitable marker of differentiation commitment. C/EBPB levels increase in all cells that are subjected to the DMI adipogenic stimulus, but the levels are not predictive of whether or not a cell will continue on to differentiate once the stimulus is removed (Bahrami-Nejad et al., 2018). Previous work using single-cell imaging showed that PPARG, the master transcriptional regulator of fat cell differentiation, exhibits bimodal, irreversible behavior and suggested that the level of PPARG can distinguish undifferentiated from differentiated cells (Ahrends et al., 2014; Bahrami-Nejad et al., 2018; Park et al., 2012) (Figures 1A-C).

**Figure 1.**
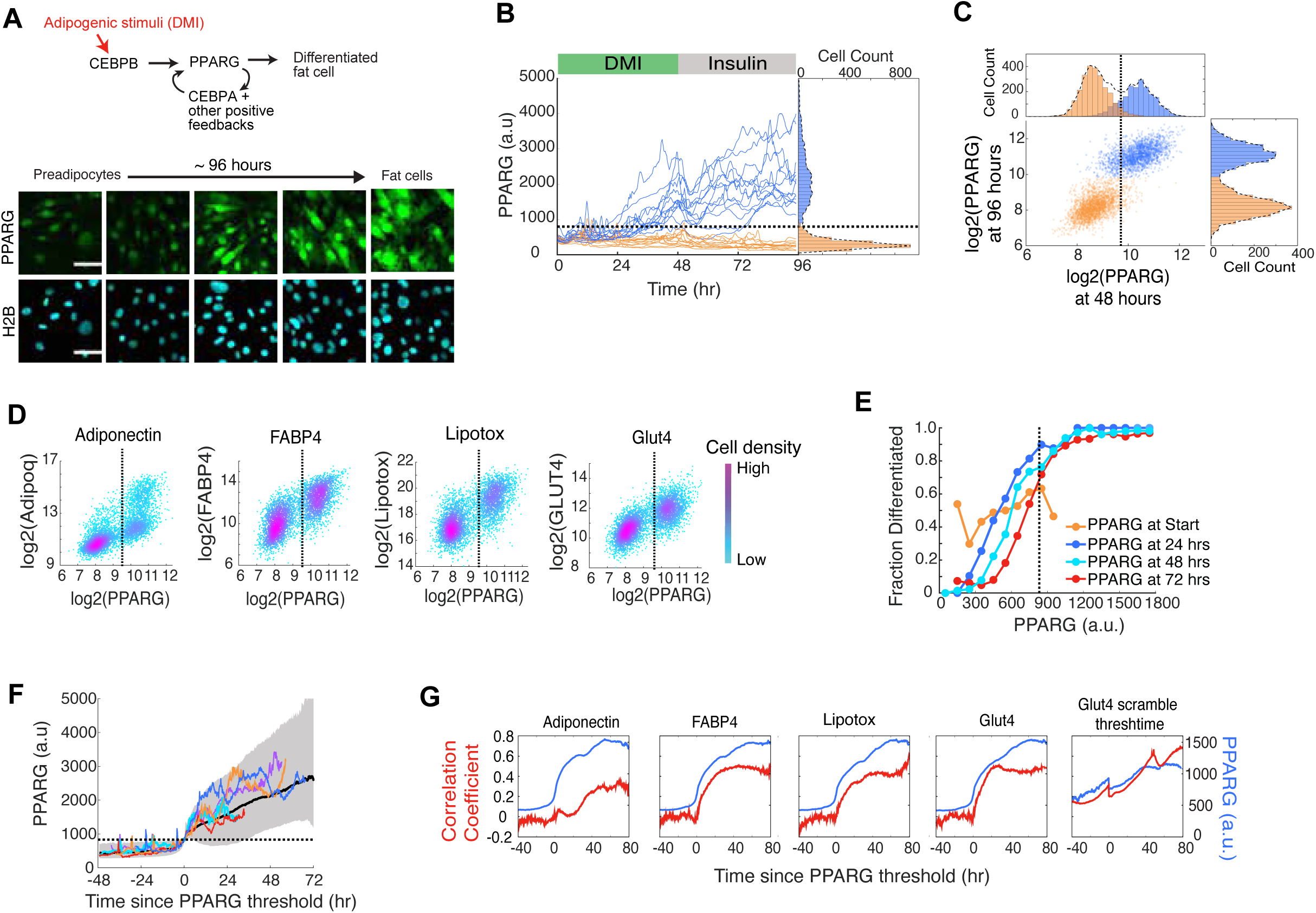
Live-cell analysis of endogenous PPARG expression shows that there is a precise time when cells irreversibly commit to differentiate. (A) Images of cells expressing endogenous citrine-PPARG that were differentiated using the standard 96-hour DMI protocol. Scale bar, 50 μm. (B) Results of a typical experiment in which thousands of single cells were stimulated to differentiate. Thirty single-cell traces are shown as examples. Representative of 4 independent experiments. (C) Scatter plot using data from (b) showing PPARG levels for each cell at 48 hours, just before the DMI stimulus was removed, and at 96 hours. Dashed black line represents the PPARG threshold (see Methods). Cells were defined as differentiated (blue) or undifferentiated (orange) by whether their PPARG levels were in the high or low intensity distributions, respectively. (D) Citrine-PPARG cells were differentiated using the DMI protocol, and immunocytochemistry was performed at 96 hours for adipocyte markers (for each scatter plot: n > 4000 cells, representative of 2 independent experiments). (E) The timecourses from (B) were split into equal-width bins by their PPARG values at 0, 24, 48, and 72 hours. The fraction of differentiated cells represents the number cells that crossed the PPARG threshold at the end of the experiment divided by the number of cells in the bin. (F) Differentiating cells from (B) were computationally aligned so that the zero timepoint represents the time when the cell crossed the PPARG threshold. We plotted 5 representative single cell traces, the median (solid black line), and the 5th-95th percentile (shaded region). (G) PPARG timecourses from the cells that differentiated after 96 hours in (D) were computationally aligned as in (F) and plotted (blue curves). At each aligned time point, the Pearson correlation coefficient between the aligned PPARG values and the endpoint immunofluorescence values for adipocyte markers was calculated (red curves). As a comparison, PPARG values for the Glut4 panel were aligned to a randomized PPARG threshold crossing point. The randomized crossing point was generated by scrambling the vector of measured threshold points for each cell so that each threshold point is matched with different cell. (B-F) The dotted black line represents the calculated PPARG threshold for that experiment.

Here we seek to understand the molecular mechanism when and how cells coordinate the commitment to terminally differentiate and exit the cell cycle, and how this coordinated process controls the number of terminally differentiated cells produced. We use adipogenesis as a model for terminal cell differentiation because it is an experimentally-accessible system in which the cell cycle and terminal differentition have been shown to be linked both in vitro and in vivo (Jeffery et al., 2015; Tang et al., 2003). We start by validating that a threshold level of fluorescently-tagged endogenous PPARG protein can be used in live cells to mark the precise time when preadipocytes irreversibly commit to terminally differentiate. By combining this live-cell PPARG sensor with a reporter to mark the G1 phase (Sakaue-Sawano et al., 2008), we establish a method that can simultaneously track both cell-cycle progression and the precise commitment point to terminally differentiate. Our live-cell measurements show that the S/G2/M phase of the cell cycle suppresses the gradual increase in PPARG expression that is triggered by addition of adipogenic differentiation stimuli. Markedly, we show that cells commit to terminally differentiate exclusively in G1 by triggering a PPARG-driven switch, and this same switch also triggers permanent cell-cycle exit by rapidly inducing high expression and increased stability of the CDK inhibitor p21. Thus, cells become post-mitotic precisely when they commit to terminally differentiate. Most importantly, we show that cells undergo a competition during each G1 phase between cyclin D1 and PPARG-induced p21 that controls whether progenitor cells have one or more cell cycles before they terminally differentiate. In this way, the competition decides whether a cell terminally differentiates and permanently stops future cell divisions, or, alternatively, starts the next cell cycle and undergoes at least one more cell division - which allows the same progenitor cell to produce more terminally differentiated cells. We show that this competition in G1 phase allows for a dual control whereby the strength of adipogenic and mitogen stimuli regulates both the number of terminally differentiated cells produced and also allows for sufficient progenitor cells to be maintained. Thus, G1 competition represents a control principle that can explain how a tissue can regulate the number of terminally differentiated cells produced while maintaining pools of progenitor cells at similar levels.

## RESULTS

### Live-cell analysis of the precise time when preadipocytes commit to terminally differentiate

A major limiting factor in understanding the relationship between the cell cycle and terminal differentiation has been the lack of a quantitative live-cell readout that can mark the precise time point when a cell commits to terminally differentiate (Buttitta and Edgar, 2007). We thus started by establishing such a live-cell readout. We chose adipogenesis as a model system for terminal differentiation since the cell cycle is known to regulate adipogenesis and the validity of using in vitro cell models for adipogenesis studies has been corroborated by in vivo studies (Ghaben and Scherer, 2019; Jeffery et al., 2015; Tang et al., 2003). Adipogenesis is centered on a master transcriptional regulator, PPARG, whose expression is driven by both external input signals and internal positive feedback loops (Ahrends et al., 2014; Rosen and Spiegelman, 2014) (Figure 1A, top). In previous work, we used CRISPR-mediated genome editing to fluorescently tag endogenous PPARG as a live-cell readout of differentiation progression (Bahrami-Nejad et al., 2018). To enable automated nuclear tracking of moving cells, we also stably transfected the cells with fluorescently labeled histone H2B (Figure 1A, bottom). We now go on to determine whether fluorescently-tagged endogenous PPARG can be used to measure a threshold level of PPARG expression that marks in live cells the precise time point when a cell irreversibly commits to terminally differentiate.

To induce differentiation, we applied a commonly used adipogenic hormone cocktail (DMI, see Methods) that mimics glucocorticoids and GPCR-signals that raise cAMP. To determine whether there is a threshold for terminal differentiation, it is critical to remove the differentiation stimulus at an intermediate timepoint to be able to determine whether or not a cell can continue on to reach and maintain a distinct terminally differentiated state days later. In our protocol, we applied the differentiation stimulus, DMI, to preadipocytes for 48 hours and then removed it, replacing it with growth medium containing insulin for an additional 48 hours at which point the differentiation state of the cells was assessed (Figure 1B). Indeed, this protocol showed two distinct outcomes for cells at 96 hours: one group of cells that kept increasing PPARG levels after removal of the differentiation stimulus (Figure 1B, blue traces), as well as a second group of cells in which the PPARG levels in cells fell back to the undifferentiated progenitor state (Figure 1B, orange traces). A histogram of the cells at 96 hours showed that the distribution of PPARG levels is bimodal with a high peak representing the differentiated cells and a low peak representing the undifferentiated cells (Figure 1B, right-side histogram).

The bimodality in the histogram of PPARG levels at 96 hours suggested that a threshold level of PPARG may exist that can determine differentiation outcome. However, to validate that there is indeed a defined PPARG threshold that can predetermine the fate of a cell days later, one needs live cell imaging. The fate of each cell is apparent from the PPARG levels at 96 hours (Figure 1B), but one needs to be able to track each cell back to before the stimulus was removed and assess whether its PPARG level before the stimulus was removed could indeed predict its final fate. Because we had timecourses of PPARG expression for each cell, we could carry out this analysis. We thus could compare for each cell its PPARG levels at 48 hours - before DMI was removed - to its PPARG level at 96 hours (Figure 1C). Indeed, the level of PPARG before stimulus removal at 48 hours could predict with a less than 5% false positive rate whether a cell will keep increasing PPARG and terminally differentiate or will lower its PPARG levels and fall back to being a progenitor cell. We could thus calculate a threshold which is determined as the center between the two peaks in the histogram at the 48-hour timepoint (Figures 1B-C, black dashed line; See Methods).

In additional control experiments, we validated that high and low PPARG expression in individual cells at 96 hours is correlated with high and low expression of different well-established markers of mature adipocytes, confirming that high PPARG is indeed a marker for the differentiated adipocyte state (Figures 1D and S1). We also confirmed that PPARG levels are predictable of final fate throughout the differentiation process, independently of when cells pass the threshold for terminal differentiation. As shown in Figure 1E, even at different timepoints throughout adipogenesis (24, 48, 72 hours), cells with higher PPARG levels have a higher probability to differentiate. Nevertheless, the probability to differentiate could not be predicted by PPARG levels at the start of the experiment (Figure 1E), suggesting that the terminal differentiation fate is not predetermined.

The analysis thus far confirmed the existence of a threshold in PPARG levels that can precisely measure the time point when cells commit to terminally differentiate during adipogenesis. But what drives cells to pass this threshold? Previous work showed that a positive feedback-driven bistable switch mechanism between PPARG and several co-regulators can amplify PPARG expression (Ahrends et al., 2014; Park et al., 2012; Wu et al., 1999). To determine whether such a positive feedback-driven bistable switch is responsible for the here identified PPARG threshold, we computationally aligned single-cell traces to the time when each cell crosses the PPARG threshold. Markedly, the aligned timecourses show a sharp sigmoidal increase from a slow rate of PPARG increase before the PPARG threshold to a fast rate after that time point. This observed switch from low to high PPARG levels at the timepoint at which the PPARG threshold is reached argues that the PPARG threshold marks the precise time when the bistable PPARG switch mechanism is triggered (Figures 1F and S1).

Lastly, we found that PPARG levels were not correlated with endpoint measurements of adipocyte markers early in adipogenesis, but once the threshold was reached, PPARG levels sharply switched to being positively correlated (Figure 1G), supporting the conclusion that crossing the PPARG threshold marks a short time window of PPARG self-amplification that causes an irreversible commitment to the future terminally differentiated adipocyte state (see also Figure S1D). However, if we had not been able to calculate and define a PPARG threshold and we were thus unable to align the timecourses, we would only see that there is an increased probabilistic relationship between PPARG expression and mature adipocyte markers; for example, see rightmost plot in Figure 1G which shows a gradual increase in correlation with PPARG with GLUT4 when timecourses are not aligned by the threshold. Thus, without being able to measure a threshold for each cell and being able to align the PPARG timecourse for each cell by this threshold, we would be unable to mark a precise timepoint for differentiation commitment, as can be seen when comparing the aligned and unaligned plots in Figures 1G and S1E.

Taken together, these different experiments validate that a threshold level can be used to mark a precise time when progenitor cells commit to terminally differentiate even before the markers of mature fat cells can be measured. When adipogenic stimuli are removed, cells that passed the PPARG threshold go on to terminally differentiate two days later, while cells below the threshold return to the undifferentiated progenitor state. Thus, fluorescently-tagged endogenous PPARG can be used to directly address the questions when a cell commits to terminally differentiate and what the connection is between the commitment to terminally differentiate and permanent exit from the cell cycle.

### Simultaneous single-cell analysis shows that further entry into the cell cycle is blocked once a cell reaches in G1 the differentiation commitment point

In order to monitor cell cycle progression and the commitment to terminally differentiate simultaneously in the same cells, we made a dual-reporter cell line by transfecting a FUCCI cell cycle reporter (Sakaue-Sawano et al., 2008) into an OP9 preadipocyte cell line we had previously generated that expressed endogenous citrine-PPARG (Figure 2A)(Bahrami-Nejad et al., 2018). The cell cycle reporter is composed of a red fluorescent protein mCherry fused to a fragment of the geminin protein that includes a degron for APC/C and is degraded by both anaphase promoting complex/cyclosomes (APC/C) (here referred to as APC/C degron reporter). Specifically, the APC/C-degron reporter signal rapidly drops when cells activate the first E3 ubiquitin ligase APC/C^CDC20^ in mitosis before cells start G1 phase, and the signal only starts to increase close to the end of G1 phase when the second APC/C^CDH1^, which is active during G1, is rapidly inactivated (Cappell et al., 2016). To validate that this APC/C reporter is suitable to monitor G1 length in the OP9 preadipocyte cell system, we compared its dynamics to that of a reporter of CRL4-Cdt2-mediated degradation that provides a more precise measure of the G1/S transition (Grant et al., 2018; Sakaue-Sawano et al., 2017). We confirmed that the expression of the APC/C degron reporter and degradation of the CRL4-Cdt2 reporter coincide in these cells (Figure S2). Thus, the dual-reporter system can be used to accurately measure the start of G1, as well as G1 length, simultaneously along with measuring the time of commitment to the terminally differentiated state. In the following experiments, we purposely used sub-confluent cell plating conditions in order to maximize the number of cell divisions, to reduce the effect of cell density on cell cycle arrest, and to improve the fidelity of the automated tracking algorithm (Figure S3A-S3B).

**Figure 2.**
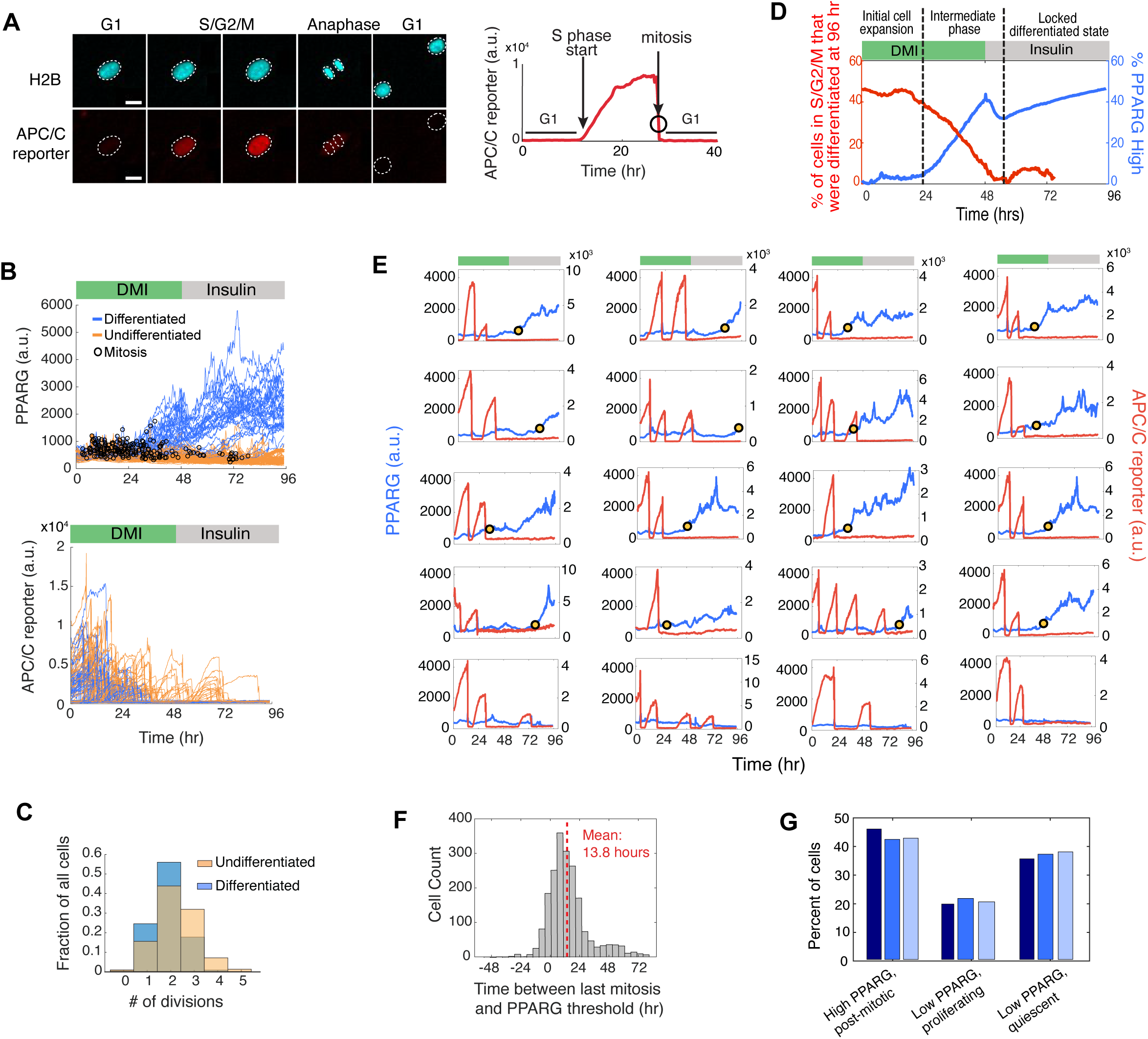
Cells commit to terminally differentiate exclusively during G1 phase. (A) Dual reporter cells were made by stably expressing H2B-mTurquoise(CFP) and APC/C-reporter-mCherry(RFP). Data from a representative single cell entering the cell cycle is shown. Anaphase is shown both by a split in the H2B-mTurquoise signal (top images) and by a sharp drop in APC/C reporter signal (bottom timecourse). Scale bar, 20 µm. White outlines mark the position of nuclei in cells after anaphase. Figure S2 confirms that the APC/C reporter is suitable to monitor G1 length in the OP9 preadipocyte cell system by showing very similar dynamics to that of a reporter of CRL4-Cdt2-mediated degradation (Grant et al., 2018; Sakaue-Sawano et al., 2017). (B) The dual reporter cells allow simultaneous measurement in thousands of single cells of differentiation state using PPARG levels (left) and cell cycle state using the APC/C sensor (right). The timepoints at which mitosis occurred were determined by using the split in H2B signal (black open circles). Cells were stimulated to differentiate with DMI using the standard adipogenic protocol. Representative of 4 independent experiments. (C) Comparison of the number of observed mitotic events that occurred in cells that were differentiated versus cells that remained undifferentiate at the end of the 96-hour experiment shown in (B). (D) Plot showing how the fraction of cells in S/G2/M (red) or with PPARG levels higher than the threshold (blue) varies during a 96-hour differentiation timecourse induced by DMI. (E) Examples of PPARG (blue) and APC/C reporter (red) timecourses obtained in the same single cell. The yellow dot in each plot marks the time at which that cell reached the PPARG threshold and irreversibly committed to the differentiated state. Bottom row shows examples of 3 undiffentiated/proliferating cells and 1 undifferentiated/quiescent cell that no longer proliferates even after a serum refresh at 48 hours. (F) Histogram of the difference between the time when the PPARG threshold is crossed and when mitosis last occurred for each cell in the experiment shown in (B). The PPARG threshold is reached on average ∼14 hours after the last mitosis is completed. Median value is 11 hours. Negative values indicate cells that reached the PPARG threshold before the last mitosis was completed. (G) Percent of differentiated/post-mitotic, undifferentiated/proliferating, and undifferentiated/quiescent cells generated in three independent DMI-induced differentiation experiments.

To determine when terminal cell differentiation occurs relative to the last cell cycle, we tracked PPARG expression and APC/C reporter timecourses over four days of differentiation. Trajectories of cells in the population that end up terminally differentiated are marked in blue, and cells that remained undifferentiated are marked in orange (Figure 2B). The trajectories show that cells that will terminally differentiate had fewer cell cycles and exited the last mitosis earlier (Figures 2B-C) compared to cells that end up not undergoing terminal differentiation. Such an inverse relationship between proliferation and terminal differentiation can be represented in a cumulative plot comparing the percent of cells still in S/G2/M versus the percent of cells that have crossed the PPARG threshold for terminal differentiation, as a function of time after DMI stimulation (Figure 2D). Control experiments showed no significant differences in PPARG levels between cells that underwent two versus three cell cycles which argued against the possibility that the lower differentiation observed in cycling cells was due to PPARG simply being diluted more in cells that cycle more often (Figure S3C-S3D).

When visually inspecting hundreds of single-cell traces, we found great variability in the kinetics of PPARG increases and number of cell-cycles before terminal differentiation (Figure 2E). However, no new cell-cycle entry was observed if the PPARG level in a cell increased above the threshold for terminal differentiation (marked with a yellow dot, Figure 2E), arguing that permanent cell-cycle exit is forced on cells when they reach the commitment point to terminally differentiate. We observed that PPARG levels already increased in many cells during S/G2/M phase of the cell cycle, but a large majority of differentiating cells only reached the PPARG threshold for terminal differentiation in G1 phase. Control experiments using a CDK2 activity reporter (Spencer et al., 2013) instead of the APC/C sensor confirmed that commitment to terminally differentiate happened during G1 phase, as shown by the fact that PPARG levels are only high when CDK2 activity levels are low (Figure S3E-S3F). Perhaps the clearest relationship between terminal differentiation and the last mitosis can be seen in a histogram analysis of when relative to the last mitosis each cell commits to a terminally differentiated state. While cells passed the commitment point for terminal differentiation at different times, almost all did so only after mitosis and the average time to commitment is approximately 14 hours from the last mitosis (Figure 2F). Thus, cells commit to the terminally differentiated state almost exclusively in G1. It should be noted that our live dual-reporter method in which we can measure cell cycle and differentiation progression simultaneously allows us to distinguish between cells that become 1) post-mitotic, differentiated; 2) undifferentiated, proliferating; or 3) undifferentiated, quiescent. For example, the bottom right plot in Figure 2E shows a cell that remains undifferentiated but becomes quiescent and ceases to proliferate even when serum is refreshed at 48 hours. The percentage of cells that end up in the three different cell fates in a typical DMI-induced differentiation experiment is shown in Figure 2G.

An interesting result from this analysis was that preadipocytes undergo a variable number of cell divisions before they differentiate (Figures 2C and 2E), arguing that terminal differentiation of adipocytes does not occur after a fixed number of cell divisions before differentiation as has been previously suggested (Tang et al., 2003). Since the previous study relied on averaged, population-based measurements, the variable number of mitoses in different cells could likely not be resolved without live single-cell analysis. Not only do the number of cell cycles vary, but there is also great variability in the time after stimulation when cells start to increase PPARG levels, and also in the time cells spend in G1 before cells reach the PPARG threshold for terminal differentiation (see also Figure S4).

We conclude that terminal adipocyte differentiation occurs after a variable rather than fixed number of cell cycles, that nearly all preadipocytes reach the commitment point for terminal differentiation after spending variable times in G1, that cells partially suppress the increase in PPARG during S/G2/M, and that cells permanently exit the cell cycle at precisely the same time when they pass the commitment point for terminal differentiation.

### PPARG regulates terminal cell-cycle exit by inducing p21 and FKBPL

The dual reporter timecourse data showed that DMI-stimulated cells that stopped proliferating earlier had also consistently higher levels of PPARG (Figures 2B, 2D, and 2E), supporting that PPARG may suppress the cell cycle and also regulate permanent cell cycle exit. To test for a direct role of PPARG in suppressing proliferation, we carried out siRNA experiments in our dual reporter cells and showed that depletion of PPARG indeed resulted in an increase of the percent of proliferating cells at all timepoints throughout the differentiation process (Figure 3A). Based on our observation that differentiation commitment occurs almost exclusively only in G1-phase and out of a state with low CDK2 activity (Figure S3E-S3F), we hypothesized that PPARG may increase the expression of one of the CDK inhibitors, which may then slow or inhibit entry into the next cell cycle.

**Figure 3.**
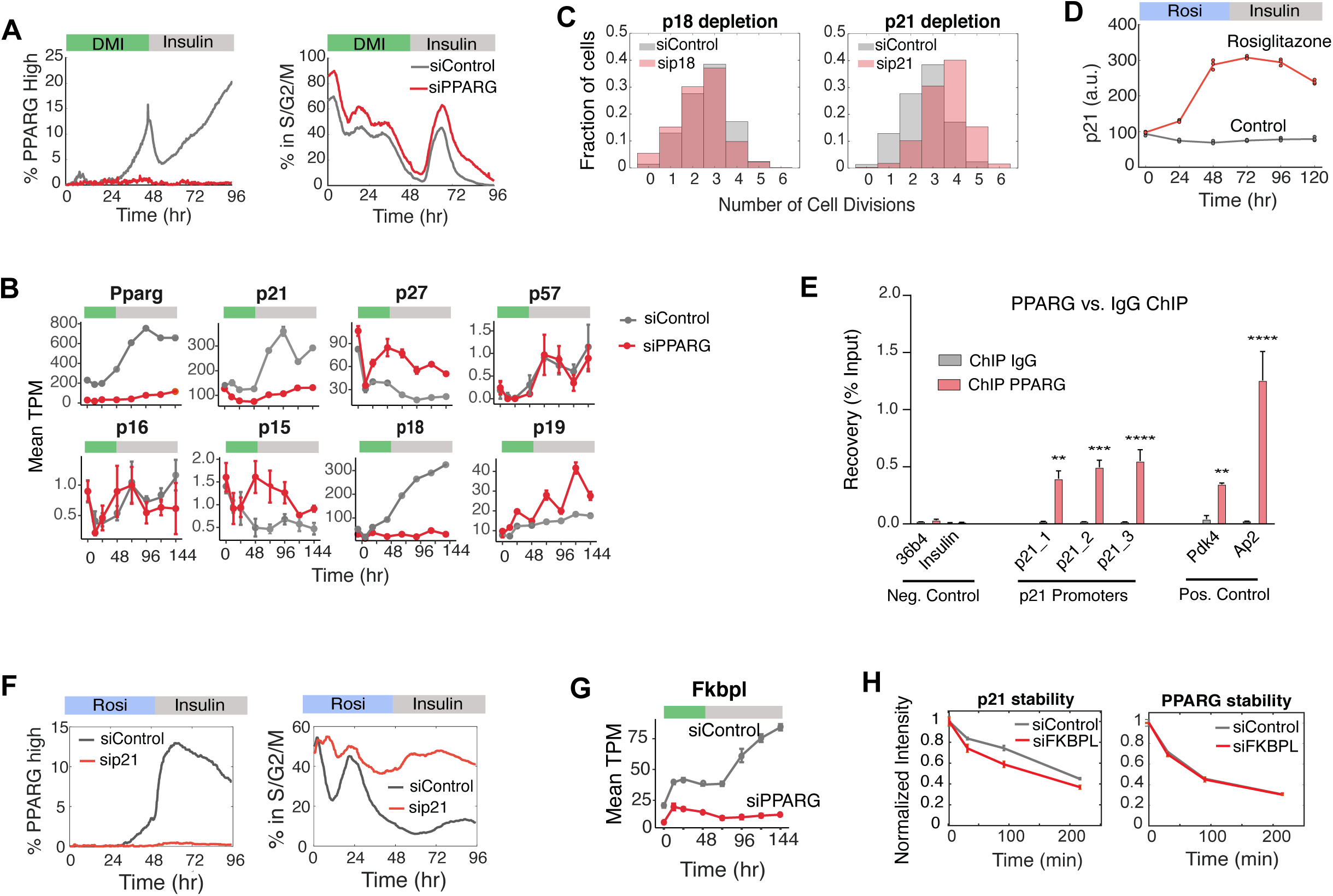
PPARG directly upregulates expression of the CDK inhibitor p21. (A) Dual-reporter cells were transfected with PPARG or control siRNA and then stimulated to differentiate. The percent of cells in S/G2/M phases at each time point is calculated by counting the cells that expressed the APC/C reporter during the 96-hour differentiation period divided by the total number of cells. The percent of PPARG high cells was assessed by counting the cells with PPARG levels above the threshold divided by the total number of cells at the respective timepoint. Cells were induced to differentiate with the standard DMI protocol. Approximately 5000 cells were analyzed per experiment. Representative of 3 independent experiments. (B) Wildtype OP9 cells were transfected with PPARG or nontargeting siRNA and then stimulated to differentiate with DMI. RNA samples were collected every 24 hours for 144 hours. Data is plotted as transcripts per million (TPM), and mean ± 1 SD is shown for three replicates. (C) Dual-reporter cells were transfected with p21, p18, or nontargeting siRNAs and stimulated to differentiate with DMI. The number of cell divisions per cell is reported in the normalized histograms. Representative of 2 independent experiments. (D) Wildtype OP9 cells were stimulated to differentiate by addition of 1 µM rosiglitazone. p21 levels were measured by fixing and staining cells at subsequent times. Approximately 5000 cells were analyzed per experiment. Data is plotted as mean (line) along with the values for each of the three experiments (points). (E) Wildtype OP9 cells were stimulated with 1 µM rosiglitazone for 48 hours. Chromatin immunoprecipitation (ChIP) of PPARG was performed followed by qPCR. Three sites on the p21 promoter are shown. The promoters of insulin and Arbp/36b4 served as negative controls, and known PPARG target genes Fabp4/aP2 and Pdk4 (pyruvate dehydrogenase kinase, isoenzyme 4) were used as positive controls. Data are normalized to a nontarget genomic site and IgG enrichment. Two biological experiments were used. Two-way ANOVA with Bonferroni’s multiple comparisons test was applied for statistical analysis. Values represent means ± SEM. p<0.05, ** p<0.01, ***p<0.001, ****p<0.0001. (F) Wildtype OP9 cells were transfected with p21 or Control siRNA and stimulated to differentiate by addition of 1 µM rosiglitazone. Differentiation and cell cycle progression were assessed in the same manner as in Figure 3A. (G) FKBPL expression under nontargeting vs PPARG knockdown were obtained from the RNA-seq data in (B). Data is reported as TPM, mean ± 1 SD. (H) Wildtype OP9 cells were transfected with FKBPL or nontargeting siRNAsand stimulated to differentiate with DMI. Stability of p21 and PPARG were assessed by adding 30 µM cycloheximide to the media 24 hours after DMI addition and then fixing and staining for protein levels at different subsequent times. Approximately 5000 cells were analyzed per experiment. Data is plotted as mean ± 1 SD of three replicates.

We sought to identify putative inhibitors of proliferation by performing comparative RNA-seq analysis using cells transfected with siRNA targeting PPARG, or control siRNA. We collected the transcripts at different timepoints during a 144-hour DMI differentiation protocol. When we examining mRNA expression profiles of canonical CDK inhibitors, we identified two that were strongly regulated by PPARG expression, p18 and p21 (Figure 3B). A PPARG-mediated increase in p21 has also been reported in other cell types (Han et al., 2004). To determine whether p18 and p21 mediate cell-cycle arrest during adipogenesis, we carried out siRNA knockdown experiments and found that p21, but not p18, knockdown led to an increase in proliferation (Figure 3C). Experiments in which p21 was knocked down with siRNA further showed that p21 is required for PPARG to mediate both terminal cell differentiation as well as suppression of proliferation (Figure 3D). We tested whether PPARG could regulate p21 expression directly by performing ChiP-Seq experiments which revealed significant binding of PPARG to the promoter of p21 during adipogenesis induced by DMI stimulation (Figure 3E). To further test whether the effect of PPARG on p21 is direct, we added rosiglitazone, a small molecule that directly activates PPARG, which led to a robust increase in p21 expression (Figure 3F).

In the same RNA-seq data, we also found that PPARG increases the expression of FKBPL (WiSP39), a protein that was shown to stabilize p21 and increase the ability of p21 to arrest cells in response to ionizing radiation (Jascur et al., 2005)(Figure 3G). To test if p21 could be stabilized by FKBPL during the early stages of adipogenesis, we carried out cycloheximide protein degradation experiments to measure the half-life of p21 in cells transfected with siRNA targeting FKBPL. Our results showed that knockdown of FKBPL causes a small decrease in p21 half-life but did not affect the half-life of PPARG, supporting that FKBPL does regulate p21 stability during terminal cell differentiation (Figure 3G). Taken together, our results demonstrate that PPARG slows G1 progression or stops cell proliferation in G1 by increasing p21 levels via increasing p21 transcription and by FKBPL-mediated slowing of p21 degradation.

### Commitment to terminally differentiate triggers immediate p21-driven cell-cycle exit

We next focused on the question how preadipocyte cells trigger permanent exit from the cell cycle once they pass the PPARG threshold for terminal differentiation. To determine the relationship between PPARG levels and terminal cell cycle exit, we took advantage of the variable increase in PPARG levels between cells in the population following DMI stimulation and grouped cells into 10 bins according to their expression level of PPARG at 48 hours (Figure 4A). At this 48-hour timepoint, the media is changed from DMI to media with insulin and growth factors and no differentiation stimuli. The corresponding mean PPARG (left) and APC/C reporter (right) signals were plotted for each bin. We found that the group of cells that passed the PPARG threshold, but not the cells that stayed below the threshold, showed no significant APC/C reporter signal in response to fresh growth media, demonstrating that cells lose the ability to re-enter the cell-cycle entry after they cross the threshold for terminal differentiation. We confirmed that this is indeed the result of reduced cell-cycle activity by calculating the fraction of cells that underwent mitosis in response to fresh growth media (Figure 4B, red). Thus, cells that pass the PPARG threshold lose their capacity to proliferate in response to freshly added growth factors, arguing that crossing the PPARG threshold marks the time when cells permanently enter a post-mitotic state.

**Figure 4.**
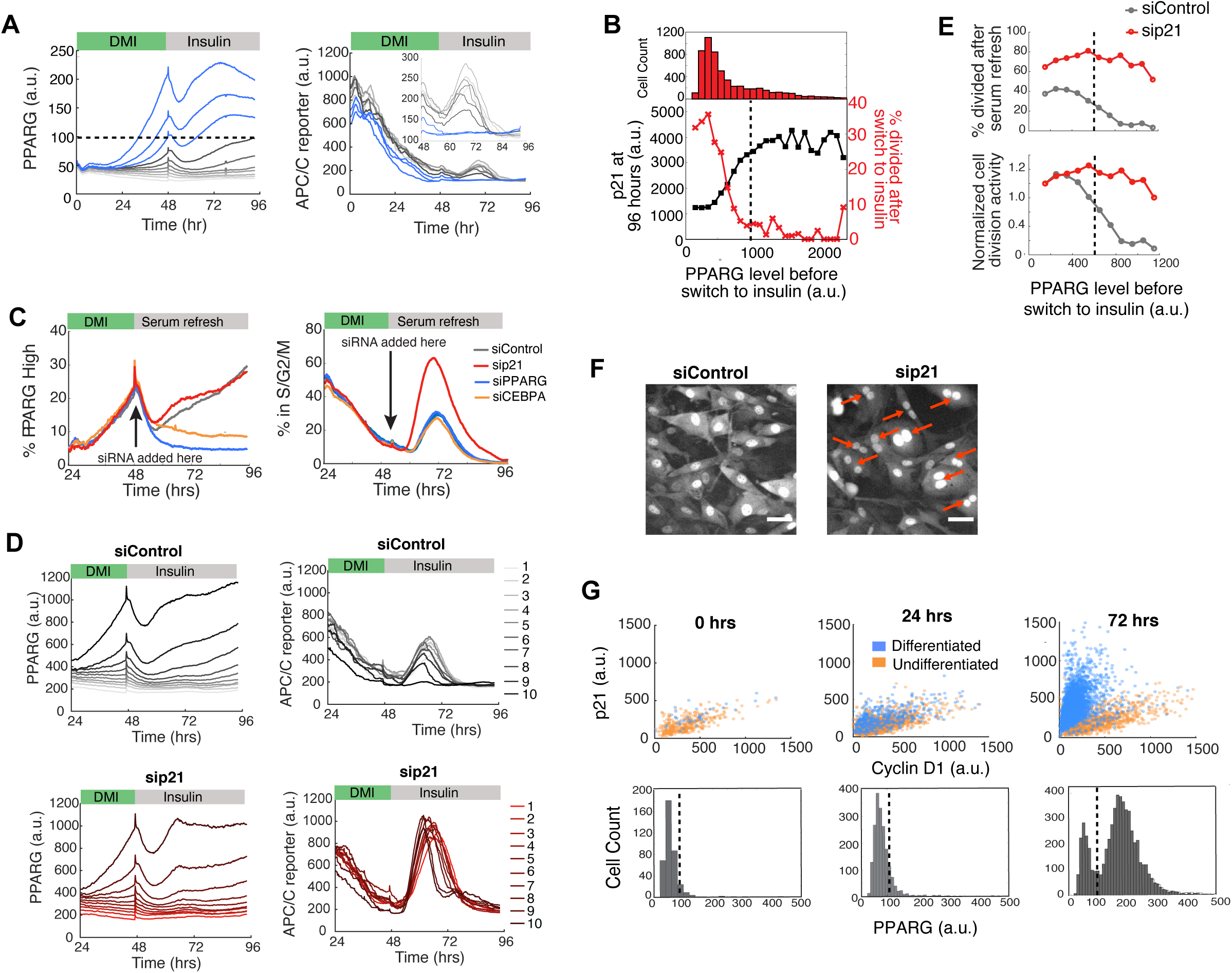
Cells simultaneously commit to differentiate and become locked in a postmitotic state by PPARG-induced maintenance of high levels of p21 expression. (A) Citrine-PPARG and APC/C reporter levels were measured in dual-reporter cells in response to media replacement at 48 hours. Cells were separated into ten bins based on their PPARG levels at 48 hours. Plotted lines show mean values for each bin. Inset shows the APC/C reporter signal between 48-96 hours. Representative of 2 independent experiments. (B) PPARG values before serum refresh were binned from 100 to 2500 a.u. in 100 a.u. increments. The plot shows for each bin the fraction of cells with a minimum of one division (measured by a split in nuclear H2B signal, red) and the final p21 level (black) and in response to serum refresh. The histogram shows the number of cells in each bin. The dotted black line shows the PPARG threshold for this experiment. Data from experiment in (A). (C) PPARG, CEBPA, p21, and nontargeting siRNA were transfected into the dual-reporter cells at 48 hours after DMI addition. siRNA knockdown efficiency is shown in Figure S6. Representative of 3 independent experiments. (D) A similar analysis as described in (A) was performed on the nontargeting and p21 knockdown conditions from (C). (E) Left, A similar analysis as in (B) was performed on the nontargeting and p21 knockout conditions from (D). Right, the same data normalized to the first PPARG bin. (F) Images of control and p21-knockdown cells from (C) obtained 48 hours after siRNA transfection (at 96 hours). Red arrows indicate representative multi-nucleated cells. Scale bar, 50 µm. (G) OP9 cells were induced to differentiate with rosiglitazone. Cyclin D1, p21 and PPARG levels were assessed by immunocytochemistry. The PPARG threshold for the whole experiment (dotted black line) was calculated at the end of the 96-hour differentiation protocol. Representative of 2 independent experiments.

We next investigated how p21 levels in individual cells change relative to PPARG levels. After completion of a live cell time course, we fixed and stained cells for p21 expression. We again binned cells according to PPARG levels and plotted the mean nuclear p21 fluorescence for each bin (Figure 4B, black). We found that p21 levels increase along with PPARG until the PPARG threshold after which p21 plateaus and stays high, suggesting that p21 is not only lengthening G1 and mediating cell-cycle exit but also maintains the postmitotic state.

To directly test for such a maintenance role of p21, we added siRNA to knockdown p21 late in adipogenesis at the 48-hour timepoint when the adipogenic stimulus was replaced with growth factor containing media. As a control, we also carried out parallel experiments in which we knocked down PPARG and CEBPA, a required co-activator of PPARG expression that is needed for cells to reach the threshold for differentiation (Bahrami-Nejad et al., 2018; Wu et al., 1999). As shown in Figure 4C, among the three regulators, only p21 knockdown resulted in a significant increase in cell-cycle activity when it was knocked down in already differentiated cells (Figure 4C). To quantitatively analyze this result, we grouped cells from the control and p21-knockdown conditions from Figure 4C by their PPARG levels at 48 hours and plotted the average time course for each group (Figure 4D). As shown in the control siRNA condition (Figure 4D, top panels), if PPARG levels are above the threshold, APC/C reporter signals remain suppressed, consistent with a lack of proliferation. However, acute knockdown of p21 expression after the differentiation commitment point had been reached (Figure 4D, bottom panels) resulted in PPARG levels no longer being able to suppress APC/C reporter signals and maintain the postmitotic state even though PPARG levels stayed high and above the threshold. Plotting the percent of cells in the cell cycle versus the level of PPARG further illustrates that if p21 is knocked down, cell divisions cannot be blocked even in cells past the differentiation threshold in which PPARG levels are high (Figure 4E). Thus, a maintained high level of p21 is required for differentiated cells to maintain the post-mitotic state. The finding that PPARG directly increases p21 levels (Figure 3), and that high PPARG levels become self-sustaining after commitment (Figure 1), explains how high levels of p21, which has a short protein and mRNA half-life of less than 30 minutes (Yang et al., 2017), can be continuously maintained to keep differentiated adipocytes permanently in a post-mitotic state.

Notably, when we examined images of cells in which p21 had been depleted late in adipogenesis after the cells had crossed the PPARG threshold, we found that the cells were enriched for multinucleation events (Figure 4F). This suggests that a critical role of p21 is to permanently prevent cell division after cells have terminally differentiated in order to prevent mitotic defects.

Finally, it was recently shown that the ratio of nuclear expression of cyclin D1 versus p21 can predict retinoblastoma (Rb) hyperphosphorylation and re-entry into the cell cycle (Yang et al., 2017). We determined whether the role of the PPARG-induced increase in p21 expression is to shift this p21-cyclin D1 ratio towards high p21 to keep Rb dephosphorylated and ensure that cells do not enter the cell cycle when mitogen stimuli increase cyclin D1 levels. As shown in a plot of p21 versus cyclin D1 levels in a large number of single cells (Figure 4G), the ratio of p21 to cyclin D1 becomes strongly skewed towards p21 when PPARG levels go above the threshold during adipogenesis, providing an explanation of how differentiated cells can maintain a robust arrested state. We conclude that a PPARG-driven rapid differentiation switch is exclusively triggered in G1 to commit preadipocytes to differentiate, and that the same differentiation switch simultaneously triggers permanent cell-cycle exit with PPARG first inducing and then maintaining p21 expression.

### An ongoing competition during G1 between terminal differentiation and continued proliferation

Previous analysis of the cell cycle in the differentiation of stem cells (Calder et al., 2012), which do not terminally differentiate, has shown that cells lengthen G1 as part of the differentiation process. The relationship between G1 lengthening and differentiation is less well understood for terminal cell differentiation, and we therefore first determined whether a gradual lengthening of G1 already occurs in the cell cycles that precedes the G1 phase from which cells terminally differentiate. Preadipocytes can undergo rapid cell cycles with a G1 phase that is on average only about 4 hours (Figure 5A). However, more than 14 hours on average is needed after the last mitosis in order for a cell to reach the PPARG threshold for terminal differentiation (Figures 2F and 5A), arguing that G1 must be lengthened before progenitor cells can commit to terminally differentiate.

**Figure 5.**
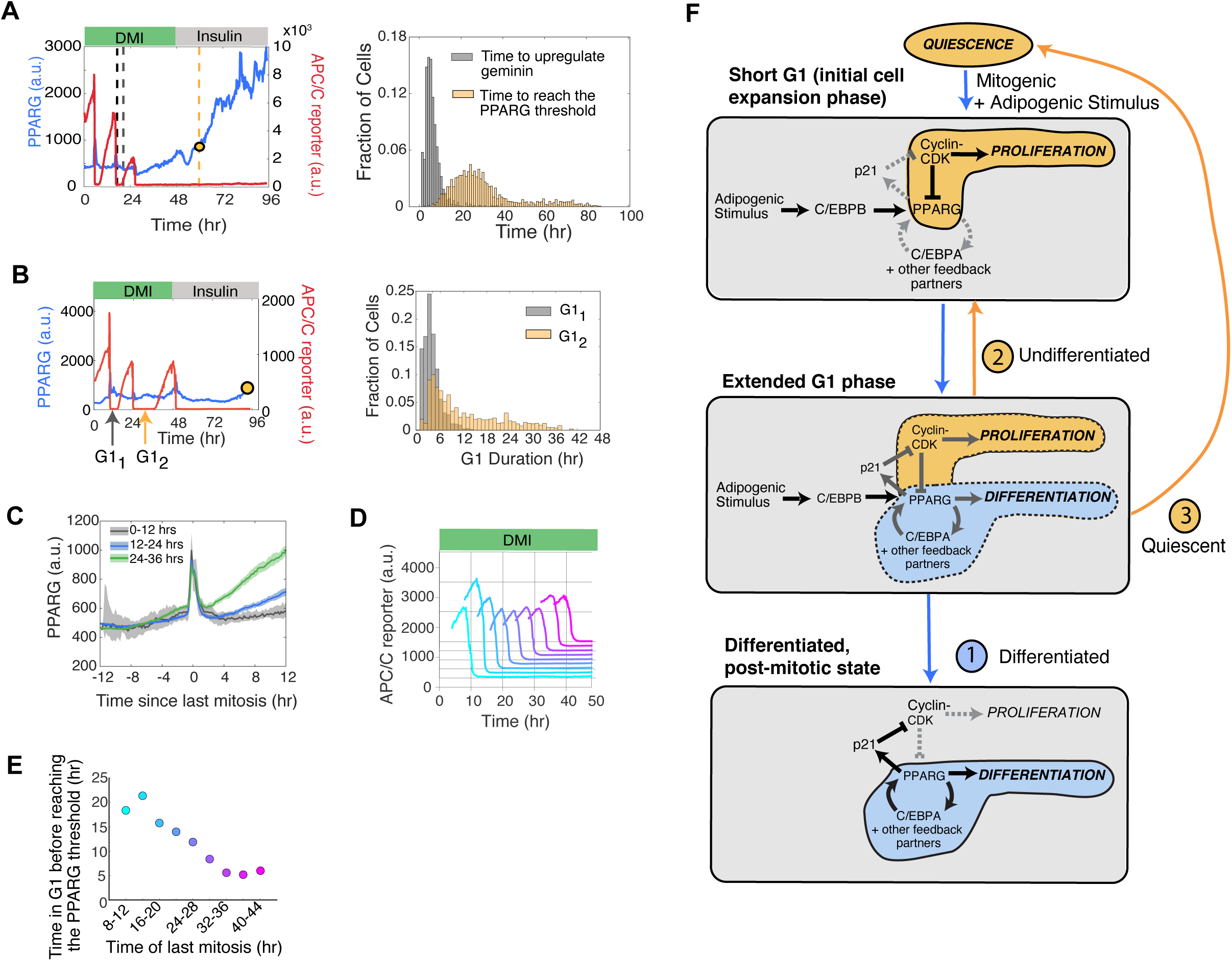
Adipogenic stimuli initiate a competition between the commitment to differentiate and entry into the next cell cycle during a gradually extending G1 phase. (A) Comparison of the time to commit to the next cell cycle (marked by upregulation of the APC/C reporter signal) versus time to commit to differentiation in cells that underwent 2-3 mitoses before differentiating. *Right*, Schematic showing that the end of the second-to-last mitosis is used as the starting reference time for each cell. *Left*, histograms comparing the two times measured in the same cell (data from Figure 2B, n > 4000 cells, representative of 4 replicates). (B) *Left*, Schematic showing which G1-periods were compared. *Right*, Histograms of the durations of the 1st and 2nd G1-periods in cells from (A) that have undergone at least 3 divisions. See also Figure S6. (C) Differentiated cells from (A) were separated into three groups based on when they last exited mitosis. The traces were aligned by the last mitosis frame. The median PPARG levels were plotted for each group (dark line). Shaded region represents the 95th confidence interval. Note that the spike in in the PPARG levels at aligned timepoint 0 is due to aligning the timecourses to mitosis. At this timepoint, there is an undefined nuclear PPARG signal because the nuclear envelope is broken down and chromatin (H2B) is condensed. (D) Timecourses from (A) were categorized into 9 bins based on time of last mitosis. The APC/C reporter peak for each bin is plotted to illustrate when the last mitosis occurred relative to addition of the adipogenic (DMI) stimulus. (E) Plot showing the average time of the last mitosis versus the average time it took for cells in that bin to increase PPARG levels to the differentiation commitment point. (F) Schematic of the three stages of G1-extension, in response to an adipogenic (DMI) stimulus. During the intermediate phase of G1-extension, stochastic competition between proliferation and differentiation causes cells to probabilistically exit into a differentiated or proliferating state. This phase defines how many differentiated cells are generated on average from a progenitor cell.

To test whether adipogenic stimuli trigger a gradual G1 lengthening over multiple cell cycles, or whether the lengthening of G1 from approximately 4 to 14 hours occurs all in a single cell before cells terminally differentiate, we selected cells that underwent three mitoses following DMI stimulation and before terminal differentiation, and compared the G1 duration for the first and second observed G1 phase (G1_1_ and G1_2_, see scheme in Figure 5B, left). Consistent with a DMI-induced gradual lengthening of G1, the second G1 length is typically significantly longer (Figure 5B, right; Figure S6A). Particularly, the duration of G1 increases strongly once the differentiation stimulus is applied for 24 hours or longer (Figure S6B). This 24-hour timepoint is also when positive feedback loops to PPARG typically start to engage (Bahrami-Nejad et al., 2018), supporting that an important role of adipogenic stimuli is to generate the G1 lengthening. Thus, adipogenic stimuli progressively lengthen G1 duration over multiple cell cycles before cells terminally differentiate out of the last G1 phase.

This DMI-induced G1 lengthening raises the question whether the reason cells do not differentiate more quickly after DMI stimulation is simply because G1 duration is so short for the first few cell cycles that PPARG can be kept mostly suppressed during S/G2/M. To test whether there is also mechanism for delaying PPARG increases that is independent from the lengthening of G1, we made use of the high variability in cell cycle responses in the cell population. By computationally aligning the timecourses by the time when the respective cell completed its last mitosis, we were able to more precisely compare PPARG increases before and after the last mitosis (Figure 5C). If a cell had its last mitosis between 0-12 hours after DMI addition, PPARG did not noticeably increase before or after mitosis. However, a small increase in PPARG could be seen after mitosis if a cell had its last mitosis 12-24 hours after DMI addition, and a very strong increase in PPARG could be observed after mitosis if a cell had its last mitosis between 24-36 hours after DMI addition. Taken together, our data supports that there is a delay mechanism that is independent of the cell cycle that restricts the increase in PPARG not only to the G1 phase, but also to a time window starting approximately 24-36 hours after DMI stimulation.

To more directly evaluate the delay before PPARG can increase in G1, we made the same alignments but with smaller time windows and by measuring the time each cell takes after mitosis to reach the PPARG threshold. We used the timecourse data shown in Figure 2B and binned cells into groups based on when a cell completed its last mitosis in increments of 4 hours (see scheme in Figure 5D). The data shows that the later a cell exited the last mitosis, and thus the longer the proliferating cell was exposed to the adipogenic stimuli, the less time a cell needs to spend in G1 before reaching the PPARG threshold (Figure 5E).

Taken together, our data can be summarized in the following model for terminal cell differentiation of adipocytes (Figure 5F): Early after adipogenic stimulation preadipocytes are proliferating and have short G1 periods. Adipogenic stimuli then gradually over time extend G1 length in sequential cell cycles and also gradually accelerate during each G1 the rate at which PPARG increases towards the threshold. The gradual lengthening of G1 and acceleration of the PPARG increase sets up a race after the end of each mitosis. In one outcome of the race, a cell first reaches the commitment to terminally differentiate, which then stops future cell cycles. In another outcome, cells end up starting the next cell cycle which allows for at least one and possibly more cell divisions before cells can terminally differentiate. There is also a third outcome that cells fail to reach the differentiation commitment point but exit the cell cycle. Thus, cells that enter G1 can either end up in a terminally differentiated state or they remain in a progenitor state where they can transition between proliferation and quiescence.

These considerations argue that there should be a competition mechanism in G1 whose role might be to balance two critical needs of a tissue, to control how many terminally differentiated cells to produce and how many progenitor cells to maintain. We hypothesized that this balance is achieved by progenitor cells delaying or accelerating the time when they commit to terminally differentiate in G1. The next series of experiments aim to test whether such a hypothesis is correct that the role of the G1 competition is to delay or accelerate the commitment point to terminally differentiate and thereby regulate a balance between the number of terminally differentiated cells produced and progenitor cells maintained.

### p21 and cyclin D1 compete to regulate the time when cells commit to differentiate and thereby control the number of terminally differentiated cells produced

Cell-cycle entry out of G1 and the length of G1 phase are not only regulated by adipogenic stimuli, as we demonstrated in Figure 5, but are also regulated by mitogens. We thus tested whether the time to reach the commitment point for terminal differentiation can also be accelerated or delayed by lowering mitogen stimuli, or by directly manipulating the regulators of G1 phase cyclin D1 and p21. We first focused on manipulating the Ras/MEK/ERK signaling pathway which is activated by most mitogen stimuli. Indeed, when we stimulated cells with DMI with and without a MEK inhibitor (Figure 6A), we observed not only that a smaller percent of cells proliferated in the presence of the MEK inhibitor but also that the time period after stimulation during which cells could proliferate was shorter. At the same time, the percentage of cells that terminally differentiated was increased and cells committed to terminally differentiate earlier. Similarly, decreasing the serum concentration along with DMI stimulation not only reduced the percent of cells that proliferated and increased the percent of terminally differentiated cells, but also cells committed to terminally differentiate earlier (Figure 6B). Thus, mitogen stimuli and the Ras/MEK/ERK pathway not only regulate the length of G1 and cell-cycle exit, but also control the time when cells commit to terminally differentiate. The most important conclusion here is that an important role of mitogens is to delay the time cells need before they reach the threshold for terminal differentiation. Since cells become post-mitotic immediately after reaching the threshold, mitogens can therefore extend the time period during which differentiating cells can proliferate and thus increase the average number of cell divisions that occur per progenitor cell before terminal differentiation.

**Figure 6.**
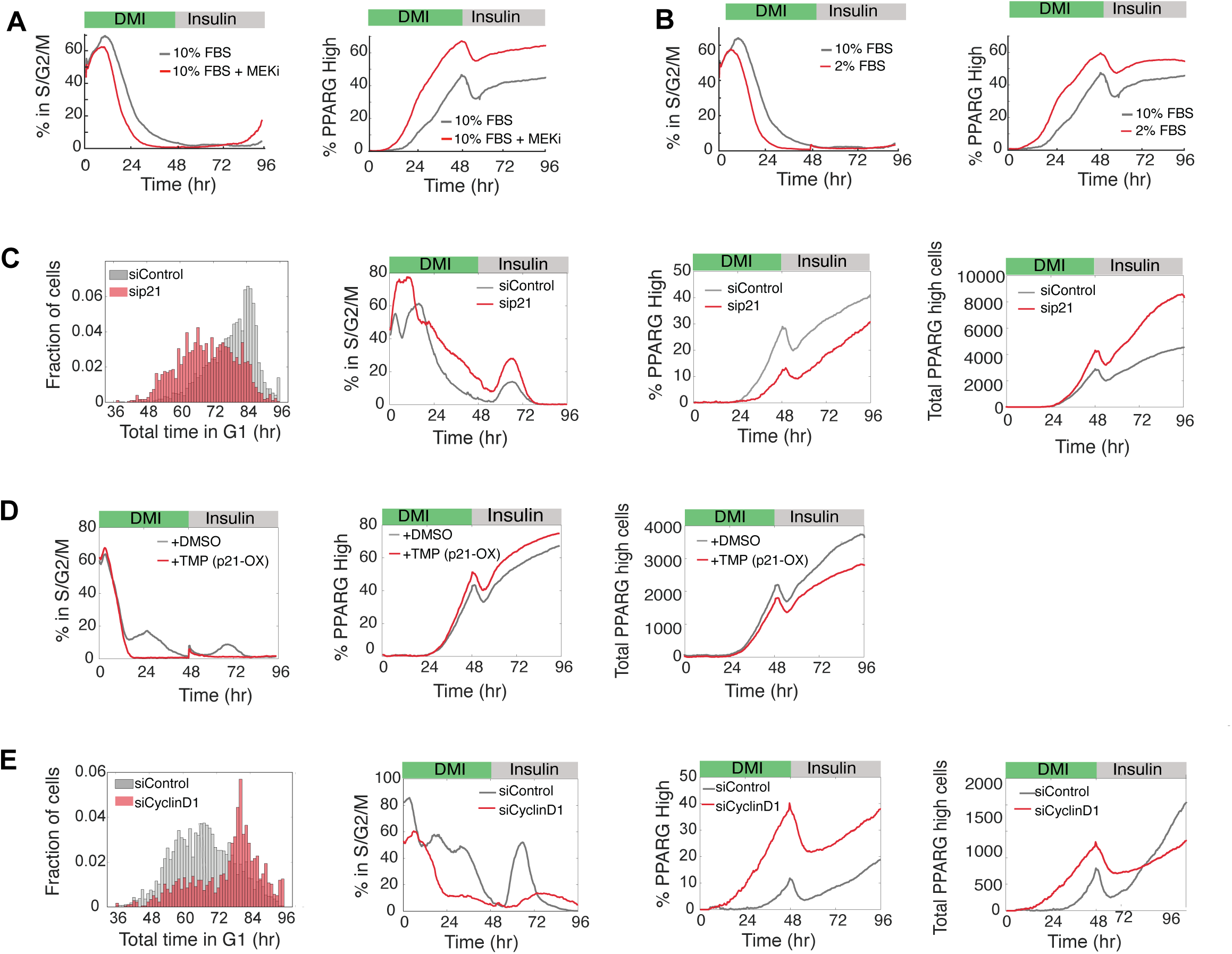
Regulation of the relative expression of p21 and cyclin D1 delays or accelerates differentiation commitment to control the total number of differentiated cells produced. (A) Dual-reporter cells were differentiated in the presence of a MEK inhibitor (PD0325091). Representative of 2 independent experiments. (B) Dual-reporter cells were differentiated in normal (10%) and reduced (2%) serum concentrations. Representative of 2 independent experiments. (C) Dual-reporter cells were transfected with p21 or nontargeting (control) siRNAs. Representative of 3 independent experiments. (D) Dual-reporter cells stably expressing a DHFR-p21-mCherry fusion protein were differentiated in the presence of 10 µM TMP (to increase expression of p21) or DMSO (control). Representative of 2 independent experiments. (E) Dual-reporter cells were transfected with cyclin D1 or nontargeting (control) siRNAs. Representative of 3 independent experiments. (A-E) Cells were induced to differentiate with the standard DMI protocol. Differentiation and cell cycle progression were assessed in the same manner as in Figure 3A. Histograms show the total time spent in G1 phase for each cell trace across all cell cycles for the respective experimental condition.

On a first glance, the results in Figures 6A and 6B support the commonly-accepted hypothesis that proliferation and terminal differentiation are opposing processes (Ruijtenberg and van den Heuvel, 2016), and one may therefore expect that mice lacking p21 or p27 should have more proliferating progenitor cells and less adipocytes. Nevertheless, one of the most striking findings from gene knockout studies was that fat tissues of female mice with deleted CDK inhibitors p21 and p27 show a synergistic 6-fold increase in the number of adipocytes (Naaz et al., 2004). We used live single-cell analysis approaches to understand this conundrum that there are more adipocytes when mice lack p21 or p27 even though mitogens have a role to suppress differentiation.

We first confirmed that cells with knocked-down p21 in vitro spend overall less time in G1 phase, consistent with p21 functioning as an inhibitor of proliferation that lengthens G1 (Figure 6C, left). Cells with knocked-down p21 also delay the commitment to terminally differentiate and can thus have more divisions before differentiation - similar to the effect we had observed with more mitogen stimulation (Figures 6A and 6B). Furthermore, we found an inverse increase in the percent of proliferating cells and a decrease in the percent of differentiated cells, which can again be explained by short G1 periods giving less opportunity for PPARG levels to increase during each G1 (Figure 6C, middle). Strikingly, however, despite the lower percentage of terminally differentiated cells, we found that the total number of differentiated adipocytes significantly increased in the p21 knockdown condition (Figure 6C, right). This increase in the total number of terminally differentiated cells can be explained by delayed commitment to terminally differentiate and a corresponding increase in the average number of cell divisions. Conversely, overexpressing p21 using a DHFR induction system yielded the opposite effect: the percent of proliferating cells decreased and there was a corresponding small increase in the percent of differentiated cells and a lower total number of differentiated cells (Figure 6D). Thus, the puzzling finding of high fat mass in mice lacking p21 or p27 type CDK inhibitors can be explained by this in vitro analysis of cells lacking p21, namely that delayed commitment to terminally differentiate results not only in a smaller percentage of cells differentiating but also in progenitor cells undergoing more cell cycles and thereby ultimately increasing the total number of produced adipocytes.

As a control, since p21 can affect both G1 and G2 phases, we carried out experiments to test whether regulation of G1 or G2 duration controls terminal cell differentiation of adipocytes. We found that the adipogenic DMI stimuli is primarily lengthening G1, rather than S/G2/M (Figure S6C), arguing that PPARG-mediated p21 expression primarily acts by lengthening G1 rather than G2-phase. Furthermore, we selectively lengthened S/G2/M by knocking down CDC25B or CDC25C, which promote CDK2/1 activation in G2, and did not observe a noticeable effect on differentiation outcome (Figure S6D-S6E).

We also tested the effects of reducing cyclin D1 expression on the total number of differentiated cells produced. As expected, we found that cyclin D1 knockdown led to an increase in G1 duration. Consistent with our model that an increase in G1 duration allows more cells to build up PPARG levels and differentiate, we observed that the cyclin D1 knockdown reduced the percent of cells that proliferated, increased the percent of terminally differentiated cells, and shortened the time for cells to terminally differentiate (Figure 6E). There were also fewer total terminally differentiated cells ultimately produced even though the percent produced was higher.

Together, these findings are consistent with the interpretation that cyclin D1 and p21 act in opposite directions in G1 to delay or accelerate terminal cell differentiation, enabling cells to regulate the number of cell divisions after adipogenic stimulation. By regulating the number of cell cycles, we hypothesized that the molecular competition mechanism in G1 has two main roles to support long-term homeostasis between progenitor and differentiated cells: To regulate not only how terminally differentiated cells are produced but also how many progenitors are maintained.

### Dual control by the strength of mitogen and differentiation stimuli can produce more or less terminally differentiated cells while maintaining similar pools of progenitor cells

Mammals maintain a large pool of preadipocytes near the vasculature in fat tissue (Tang et al., 2008; Tchoukalova et al., 2004) that is believed to be maintained by differentiation of adipose-derived stem cells and proliferation of preadipocyte progenitor cells. Yet adipocytes are replaced only at a low rate (Spalding et al., 2008), arguing that adipogenesis is a relatively slow process with only a small percentage of progenitor cells differentiating at a given time. This slow rate of differentiation motivated earlier studies that showed that the strength of adipogenic stimuli can control the percent of progenitor cells that terminally differentiate (Ahrends et al., 2014; Park et al., 2012). At the same time, the data in Figure 6 suggest that mitogen stimuli indirectly regulate the number of terminally differentiated cells as well as the number of remaining progenitor cells by regulating the number of progenitor cell divisions before cells commit to terminally differentiate.

As shown in the scheme in Figure 7A, when the commitment to terminally differentiate is delayed, extra divisions can lead to more differentiated cells. For example, when the commitment to terminally differentiate is early, a progenitor cell may have on average only one cell division and a final outcome might be one differentiated cell per activated progenitor cell. This same differentiation program has the interesting characteristic that it can, in principle, maintain a similar number of progenitor cells at the end as were present at the start. The schematic includes different outcomes of cell divisions to highlight the observed variability in the differentiation response observed between progenitor cells. In a different example, a progenitor cell with a delayed commitment may have on average 2 cell divisions and a final outcome might be three differentiated cells per progenitor cell, while again allowing that a similar number of progenitor cells is maintained during the differentiation process.

**Figure 7.**
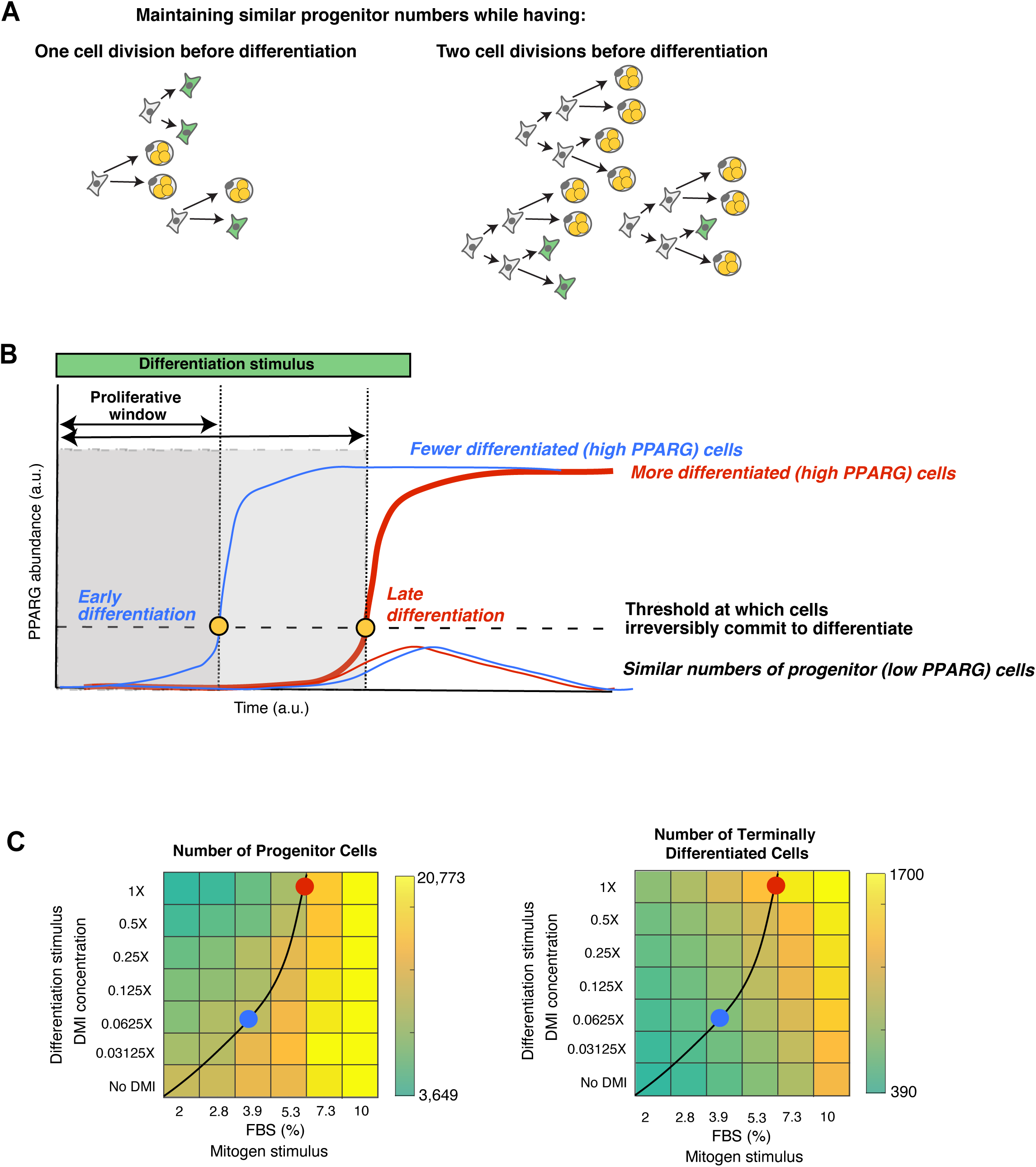
Optimal relative strengths of mitogen and differentiation stimuli can regulate the number of differentiated cells produced while maintaining similar progenitor pools. (A) Schematic showing how variable numbers of cell divisions can control the total number of terminally differentiated cells produced while maintaining similar numbers of progenitor cells. Both scenarios maintain a pool of 3 progenitor cells (green cells) while producing very different numbers of differentiated cells (3 versus 9). (B) Schematic of how delaying the time to reach the differentiation commitment threshold, and thus increasing the time when progenitor cells can proliferate before differentiating and becoming post-mitotic, increases the total number of differentiated cells. (C) Both the mitogenic and adipogenic stimuli must be simultaneously regulated in order to control the total number of differerentiated cells produced while keeping the number of progenitor cells fixed.

If a tissue needs more differentiated cells, why not just increase the strength of adipogenic stimuli or, alternatively, just increase the strength of mitogen stimuli to have more rounds of divisions before terminal differentiation? The challenge is that a tissue has to balance two needs: to produce new differentiated cells while also maintaining healthy numbers of progenitor cells to support long-term tissue homeostasis. For example, if progenitor cells are exposed to a low differentiation and high mitogen stimulus, the number of differentiated cells would increase but at the cost of ending up with a large pool of progenitor cells. If instead progenitors are exposed to a high differentiation and low mitogen stimulus, the number of differentiated cells would also increase but at the cost of depleting the pool of progenitor cells. We hypothesized that a main role of the G1 competition mechanism we identified is to facilitate a balance between differentiated cells produced and progenitor cells maintained. Figure 7B describes a simplified model how combined mitogen and differentiation stimuli may achieve this balance by employing G1 competition to regulate if and when the terminal differentiation switch is triggered and allowing for a control of the average number of cell cycles before differentiation commitment. By delaying or accelerating the commitment to terminally differentiate and regulating how many cells reach the commitment point, mitogen and differentiation stimuli may be able to regulate the number of differentiated cells produced while also maintaining similar numbers of progenitor cells.

To test the hypothesis that the strengths of both mitogen and adipogenic stimuli are critical for regulating the numbers of terminally differentiated and progenitor cells, we performed experiments in which we systematically applied different relative strengths of adipogenic and mitogenic stimuli for 48 hours and then removed the stimulus such that we could determine at 96 hours the number of terminally differentiated cells versus the number of progenitor cells. Markedly, application of increasing mitogen stimuli resulted in higher levels of progenitor cells and lower levels of differentiated cells independently of the strength of the adipogenic stimulus (Figure 7C). In contrast, the number of progenitor cells decreased and differentiated cells increased along with increasing the adipogenic stimulus independent of the mitogen stimulus. This opposite regulation argues for a control principle that it is the strength of the differentiation stimulus that triggers a fraction of progenitor cells to terminally differentiate, and that the relative strength of the differentiation versus mitogen stimulus sets up the competition in G1 that regulates the number of cell cycles. In this way, different external stimuli can control how many differentiated cells are produced and progenitor cells maintained.

Our titration experiments show that in order to increase the number of differentiated cells produced while maintain similar numbers of progenitor cells, both differentiation and mitogen stimuli have to increase or decrease in tandem. This co-dependence can best seen in the heatmaps in Figure 7C by manually plotting a contour line that connects differentiation versus mitogen stimulus conditions that maintained similar numbers of progenitor cells (black curved line in Figure 7C). The blue and red dots mark differentiation versus mitogen conditions that produce low and high numbers of terminally differentiated cells, respectively, while maintaining similar levels of progenitor cells.

## DISCUSSION

We used adipogenesis as a model system to understand how cells coordinate terminal differentiation and cell-cycle exit and developed a method to measure the precise moment of terminal differentiation while simultaneously monitoring when cells enter and exit G1 phase. With this approach, we showed that following adipogenic stimulation, progenitor cells undergo one or more cell cycles before they reach a sharp commitment point where they terminally differentiate nearly exclusively in G1. We further show that the underlying rapid PPARG-driven switch mechanism not only commits cells to terminally differentiate but also rapidly induces expression of the CDK inhibitor p21 to force permanent exit from the cell cycle.

Mechanistically, we show that the rapid increase in p21 is mediated by PPARG-induced transcription and by expression of FKBPL and a FKBPL-mediated increase in p21 half-life. The earlier finding that p21 has a short-half-life of less than an hour at both the protein and mRNA levels (Figure 3H and Yang et al., 2017) raises a question common to all terminal differentiation processes: how can a permanent post-mitotic state be maintained? In adipogenesis, PPARG can function as a continuously active driver of p21 expression in differentiated cells. Once PPARG levels reach the threshold for terminal differentiation commitment, the upregulated permanent activation of positive feedback loops self-sustain the level of PPARG independently of the input stimulus and can therefore permanently induce p21 and thus permanently prevent cells from re-entering the cell cycle. We show that acutely suppressing p21 after cells have passed the threshold for terminal differentiation re-activates the cell cycle but also results in mitotic defects, arguing that maintaining high p21 levels after the commitment to differentiate is critical for cell health. Notably, in our experiments we plated cells at low density conditions such that p21 was the dominant CDK inhibitor. At higher cell densities, the homolog CDK inhibitor p27 becomes the main regulator of CDK activity and likely has a synergistic role along with p21 in regulating terminal cell differentiation. Such a synergistic role of both CDK inhibitors is consistent with knockout data in mice which showed a 6-fold increase in fat mass when p27 and p21 where knocked out together as compared to smaller increases with only individual knockouts (Naaz et al., 2004).

Importantly, our experiments revealed that the length of G1 and number of cell divisions are regulated during the differentiation process by a competition mechanism between cyclin D1 and PPARG-induced expression of p21. This competition during G1 phase controls whether or not a cell enters the next cell cycle or terminally differentiates. At the same time, we found that PPARG expression is delayed by repression during the S/G2/M period of the cell cycle and that the self-amplification of PPARG becomes only gradually sensitized over a 36 hour-long time period following adipogenic stimulation. As a consequence of regulating these different timing mechanisms, we show that cells delay or accelerate the time when they commit to terminally differentiate which in turn controls the number of cell cycles per progenitor cells and the number of terminal differentiated cells produced on average from progenitor cells.

Finally, we show that this molecular competition during G1 can be regulated by the strength of both external adipogenic and mitogen stimuli and that increasing either adipogenic or mitogen stimuli can increase the number of terminally differentiated cells (Figures 6 and 7; see also Ahrends et al., 2014; Park et al., 2012). However, adipogenic stimuli increase the number of differentiated cells by directly depleting the existing number of progenitor cells while mitogen stimuli increase the number of differentiated cells by first generating a large pool of progenitor cells out of which only a few cells differentiate (Figure 7). This dual control mechanism can, for example, explain the conundrum how mice lacking p21, which mimics a state of higher mitogenic signaling, can have significantly higher numbers of adipocytes than control mice, despite proliferation and terminal differentiation being opposing processes (Naaz et al., 2004). Our results show that mitogen and adipogenic stimuli can coordinate a robust terminal differentiation process by controlling both how many terminally differentiated cells are produced while also ensuring that a pool of progenitor cells is maintained. Intriguingly the question of maintaining a balance of progenitor and terminally differentiated cells is particularly relevant in the context of generating complex organs such as the spinal cord in which the the different neuronal domains along the dorsal-ventral axis are formed through terminal differentiation of different cell types in a temporal sequence from the same progenitor pool (Molina and Pituello, 2017; Sagner and Briscoe, 2019). Not having enough terminally differentiated cells or an abnormally depleted progenitor pool at any stage in development could lead to deformities and disease.

Together, the opposing roles of cyclin D1 and p21 in regulating the time when cells commit to terminally differentiate (Figure 6), and the large in vivo effect of p21 and p27 knockouts on fat mass (Naaz et al., 2004), suggest that therapeutic strategies aimed at regulating the time when cells commit to terminally differentiate may proof useful to control the size of terminally differentiated tissues. For example, since DNA damage and aging increase p21 expression, such conditions may cause an imbalance between progenitor and differentiated cells and may result in insufficient fat and other terminally differentiated cells produced through this competition mechanism. It is suggestive to propose that our finding that cyclins and CDK inhibitors can act in opposite ways to control the number of terminal differentiated cells produced and progenitor cells maintained may represent an example of a more general control principle that the size of tissues may generally be controlled by a competition between cyclins on one side and drivers of terminal cell differentiation and CDK inhibitors on the other side.

## METHODS

### Generation of PPARG and APC/C dual-reporter cell line

OP9 cells with endogenously tagged citrine-PPARG2 and stably infected H2B-mTurqoise was generated as previously(Bahrami-Nejad et al., 2018). Lentivirus was generated for the APC/C reporter from the vector pCSII-EF-Geminin(1-110)-mCherry. A third-generation lentiviral packaging system was used and consisted of the following packaging vectors: pMDlg, pCMV-VSVG, and pRSV-Rev. The APC/C reporter was then stably infected into H2B-mTurqoise/citrine-PPARG2 cells to generate the dual-reporter cell lines. Selection of dual-reporter cells was done with FACS for mCherry-positive cells.

### Generation of a PPARG, APC/C, and CDK2 triple reporter cell line

Lentivirus was generated for the CDK2 sensor from the vector pCSII-EF-DHB-mTurquoise (gift from the lab of Tobias Meyer) in the same manner described above and used to infect the dual reporter PPARG/geminin cells. Selection of triple-reporter cells was done with FACS for cells that were positive for both mCherry and mTurquoise.

### Generation of a PPARG and CRL4-CDT dual reporter cell line

The CRL4-Cdt2 construct was developed by Atsushi Miyawaki’s lab (Sakaue-Sawano et al., 2017) and was obtained from the lab of Tobias Meyer. We changed the fluorescent tag to iRFP670 and generated lentivirus in the same manner described above. Selection of triple-reporter cells stably expressing iRFP670-CRL4-Cdt2 was done with FACS for cells that were positive for both mCherry and iRFP670.

### Cell culture and differentiation

Wildtype and reporter OP9 cell lines were cultured according to previously published protocols (Ahrends et al., 2014; Bahrami-Nejad et al., 2018; Wolins et al., 2006). Briefly, the cells were cultured in growth media consisting of MEM-α media (ThermoFisher Scientific) containing 100 units/mL Penicillin, 100mg/mL Streptomycin, and 292 mg/mL L-glutamate supplemented with 20% FBS. To induce differentiation, two methods were used. In the first method, a standard DMI cocktail was used: cells were treated with 125 µM IBMX (Sigma-Aldrich), 1 µM dexamethasone (Sigma-Aldrich), and 1.75 nM insulin (Sigma-Aldrich) for 48h, followed by 1.75 nM insulin for 48h. In the second method, cells were treated with 1 µM of Rosiglitazone (Cayman, USA) for 48 hours, followed by 1.75 nM insulin for another 48 hours. For fixed cell experiments, the differentiation stimuli were added to the growth media described above with one modification: 10% FBS was used (instead of 20% FBS) during differentiation conditions. The one exception i in the reduced serum experiments in Figure 4D, in which 2% FBS was used in the growth media during differentiation. For all live cell experiments, the differentiation stimuli were added to Fluorobrite DMEM media (ThermoFisher Scientific) containing 100 units/mL Penicillin, 100mg/mL Streptomycin, and 292 mg/mL L-glutamate supplemented with 10% FBS. For Figure 4D, EGF (Sigma-Aldrich E9644) was used at a final concentration of 1 μg/mL, and a MEK inhibitor PD0325091 was used at a final concentration of 100 nM.

### siRNA-mediated gene silencing

siRNA targeting *Pparg, Cebpa, p21, CyclinD1, Fkbpl* and the AllStars Negative Control siRNA were purchased from QIAGEN. For siRNA knockdown in the live-cell imaging experiments in dual-reporter cells (Figure 4, Figures 5a, 5d, and 5e), OP9 cells were transfected by reverse-transfection using µL Lipofectamine RNAiMax (Invitrogen). Briefly, our reverse-transfection protocol per well is as follows: mixed 20 µL of Optimem, 0.5 µL of a 10 µM siRNA stock solution, and 0.3 µL of RNAiMax. Let solution incubate at room temperature for 10 minutes and then add 80 µL of culture media containing the desired number of cells per well. Then the entire (∼100µL) volume is plated into one well of a 96-well plate. The siRNA/RNAiMax mixture was left on the cells for 24 hours before being aspirated away and replaced with fresh culture media containing DMI to begin the differentiation protocol.

For the live-cell imaging experiments in dual-reporter cells transfected at the 48-hour timepoint (Figure 6), the following protocol per well was used: siRNA mixture was prepared using 0.6 µL Lipofectamine RNAiMAX, 0.5 µL of a 10 µM siRNA stock solution, and 20 µL of Optimem. Incubate the mixture for 10 minutes then add 180 µL of Fluorobrite media consisting of 1.75 nM insulin. The entire solution (∼200µL total volume) was then added to cells at the 48-hour time point and left on until the end of the experiment.

### Overexpression of p21

A retroviral vector containing DHFR-Chy-p21(Spencer et al., 2013) (gift from the lab of Tobias Meyer) was used to generate viral particles to stably infect DHFR-Chy-p21into a modified dual-reporter cell line. This cell line was also stably infected with H2B-iRFP670 and a version of the APC/C reporter fused to mCerulean3. Positive clones were selected for by FACS in cell culture media containing 10 µM TMP. Cells were sorted into culture media with no TMP and grown in the absence of TMP. All overexpression experiments were done by adding 10 µM TMP into the culture media or differentiation media. In control experiments, 10 µM DMSO was added instead of TMP.

### Immunofluorescence (IF) staining

All cultured cells were fixed with 4% PFA in PBS for 30 min at room temperature, followed by five washes with PBS using an automated plate washer (Biotek). Cells were then permeabilized with 0.1% Triton X-100 in PBS for 15 minutes at 4°C, followed by blocking for 1 hour in 5% bovine serum albumin (BSA, Sigma Aldrich) in PBS. The cells were incubated with primary antibodies in 2% BSA in PBS overnight at 4°C: mouse anti-PPARγ (Santa Cruz Biotech, sc-7273, 1:1,000), rabbit anti-CEBPα (Santa Cruz Biotech, sc-61, 1:1,000), mouse anti-p21 (Santa Cruz Biotech, sc-6246, 1:100), cyclinD1 (Abcam, ab137145,1:1,000), adiponectin (Abcam, ab22554, 1:1,000), Glut4 (Santa Cruz Biotech, sc-1608,1:500), FABP4 (R&D Systems, AF1443, 1:1,000). After washing, cells were incubated with Hoechst (1:20,000) and secondary antibodies in 2% BSA / PBS for 1 hour. Secondary antibodies included AlexaFluor-conjugated anti-rabbit, anti-mouse, and anti-goat antibodies (Thermo Fisher Scientific). All secondary antibodies were used at a 1:1,000 dilution. Where indicated, lipids were co-stained by adding HCS LipidTOX Deep Red Neutral Lipid Stain 637/655 (1:1,000), ThermoFisher Scientific H34477) to secondary antibody solution. Cells were washed five times with PBS in an automated plate washer prior to imaging. For fixed-cell timecourse experiments, approximately 7,000 wildtype or dual-reporter OP9 cells were used to calculate mean values at each timepoint for each technical replicate.

### RNAseq

siRNA targeting Pparg (# L-040712-00-0005) and Negative Control siRNA (# D-001810-10-05) were purchased from Dharmacon and transfected into OP9 cells using Lipofectamine RNAiMax (Invitrogen) according to the manufacturer’s protocol. siRNA was used at a concentration of 25 nM, and the RNAiMAX/siRNA mixture was applied for 48 hours prior to the induction of differentiation. For gene expression analysis of OP9 cell samples, the cells were differentiated for 144 hours using a previously described protocol (Ahrends et al., 2014). RNA from three independent biological experiments were collected at different time points before and after induction of differentiation including (d0-d6) the extraction was completed using RNeasy Mini Kit (QIAGEN, Cat. 74104). RNA quality of all samples (n=7 time points and n=3 experiments from independent passages) was evaluated by both Nanodrop for (A260/280 >2) and Bioanalyzer 2100 High Sensitivity RNA Analysis chips (Agilent, Cat. 5067-1513) which displayed intact RNA integrity (RIN >9). mRNA samples were concentrated to ≤ 5 µl by MinElute column (QIAGEN, Cat. 74204). For generation of RNA-seq libraries, polyadenylated mRNA was isolated from 300 ng of total RNA by incubation with oligo-DT attached magnetic beads and followed by strand-specific library preparation using the TruSeq Stranded mRNA Library Preparation kit (Ilumina, Cat. 20020595). Briefly, isolated polyadenylated mRNA was fragmented using divalent cations under elevated temperature and 1^st^ and 2^nd^ strands DNA were synthesized using SuperScript II Reverse Transcriptase (provided with Ilumina kit). A-tailing and adapter ligation was performed according to the manufacturer’s protocol; the resulting dsDNA was enriched in a PCR reaction based on predetermined CT values and cleaned using AMPure XP beads (provided with Ilumina kit). Concentrations of enriched dsDNA fragments with specific adapters were determined and base pair average size as well as library integrity were analyzed using the Bioanalyzer DNA High Sensitivity chips (Agilent, Cat. 5067-4626). Samples were pooled and sequenced on the Illumina NextSeq 500/550 High Output platform (Illumina, FC-404-2002) up to 18 samples per lane with 1% PhiX spike as a control.

The read quality of the raw FASTQ files was checked with FastQC(Andrews and Babraham Bioinformatics, 2010) (v0.11.7). Next, reads were pseudo-aligned to the mouse reference transcriptome (Mus_musculus.GRCm38.cdna) using Kallisto(Bray et al., 2016) (v0.44.0) with the quantification algorithm enabled, the number of bootstraps set to 100, and run in paired-end mode. The Kallisto output files were read into R using Sleuth, and the transcripts per million (TPM), a measurement of the proportion of transcripts in the RNA pool, was used for downstream differential expression analysis(Pimentel et al., 2017).

### Measuring protein decay rates using cycloheximide

Protein decay rates were quantified as previously described(Bahrami-Nejad et al., 2018). Briefly 10,000 OP9 cells were seeded in 96-well plates) one plate for each timepoint. Cells were induced to differentiate with DMI for 24 hours. Cyclohexamide was added to the media at a final concentration of 30 μM. Cells were fixed and stained at different times after addition of cyclohexamide, and immunofluorescence was used to quantify protein concentration. Half-lives were obtained by fitting first order exponential decay curves to the data.

### Fluorescent imaging

Imaging was conducted using an ImageXpress MicroXL (Molecular Devices, USA) with a 10X Plan Apo 0.45 NA objective. Live fluorescent imaging was conducted at 37°C with 5% CO_2_. A camera bin of 2×2 was used for all imaging condition. Cells were plated in optically clear 96-well plates: plastic-bottom Costar plates (#3904) for fixed imaging or Ibidi µ-Plate (#89626) for live imaging. Living cells were imaged in FluoroBrite DMEM media (Invitrogen) with 10% FBS, 1% Penicillin/Streptomycin and insulin to reduce background fluorescence. Images were taken every 12 min in different fluorescent channels: CFP, YFP and/or RFP. Total light exposure time was kept less than 700 ms for each time point. Four, non-overlapping sites in each well were imaged. Cell culture media were changed at least every 48h.

### Imaging data processing

Data processing of fluorescent images was conducted in MATLAB R2016a (MathWorks). Unless stated otherwise, fluorescent imaging data were obtained by automated image segmentation, tracking and measurement using the MACKtrack package for MATLAB. Quantification of PPARG- and CEBPA-positive cells in fixed samples was based on quantification of mean fluorescence signal over nuclei. Cells were scored as PPARG- and CEBPA-positive if the marker expression level was above a preset cut-off determined by the bimodal expression at the end of the experiment.

For live imaging data of OP9 cells, the CFP channel capturing H2B-mTurqoise fluorescence was used for nuclear segmentation and cell tracking. Obtained single-cell traces were filtered to removed incomplete or mistracked traces according to the following criteria: cells absent within 6 hours of the endpoint, cell traces that started more than 4 hours after the first timepoint, cells that had large increase or decrease in PPARG intensity normalized to the previous timepoint, cells where H2B drops did not match drops in the APC/C reporter. If cells were binned according to their PPARG expression, cells were binned based on their mean nuclear PPARG expression at the described timepoints.

The percent of cells in the S/G2/M phases at each time point is calculated by counting the cells that expressed the APC/C reporter during the 96-hour differentiation period divided by the total number of cells. The percent of PPARG high cells was assessed by counting cells that above the PPARG threshold at that time point and dividing by the total number of cells at that time point.

### Estimating a differentiation commitment point (i.e. PPARG threshold)

PPARG values at the end of a differentiation experiment typically exhibit a bimodal distribution. In order to estimate a commitment point, PPARG values at the last frame of the experiment was fit to a 2 component gaussian mixture model. Cells were then classified as either differentiated or undifferentiated based on whether they more closely associated with the high or low component of the mixture model, respectively. The commitment point was then assessed as the value of PPARG at the 48-hour time point, before the stimuli was removed, that predicted the final differentiation classification with a false positive rate of less that 5%. In experiments where multiple conditions are present, the gaussian mixture model was only fitted to the negative control and the commitment point was selected based on the negative control model and applied to all other conditions in the same experiment.

### Statistics

Unless specified otherwise, data are expressed as mean +/-standard error of the mean (S.E.M). Live traces are expressed as median +/-interquartile range (25^th^-75^th^ percentiles). For histograms with a y-axis labeled “Fraction of Cells,” each histogram (not each plot) is normalized to the total number of cells in the population of that histogram such that all bars in the histogram add to 1. Representative results are representative of at least two independent experiments.

## Data availability

All relevant data from this manuscript are available upon request.

## AUTHOR CONTRIBUTIONS

M.L.Z. and M.N.T. conceived experiments. M.L.Z., K.K., A.R., and Z.B. performed experiments and analyzed data. M.L.Z. and B.T. wrote the image analysis scripts. M.L.Z. and M.N.T wrote the paper with input from all authors.

## ACKNOWLEDGEMENTS

This work was supported by National Institutes of Health RO1-DK101743, RO1-DK106241, P50-GM107615, a Stanford BioX Seed Grant, and a Stanford Diabetes Research Center Seed Grant (to M.N.T.), NIH F32 Postdoctoral Fellowship 5F32DK114981-02 (to B.T.), and T32-NIH T2HG00044 and NIH F31 Predoctoral Fellowship 1F31DK112570-01A1 (to M.L.Z.). The authors would like to thank Tobias Meyer, Takamasu Kudo, Antoine Zalc, Lindsey Pack (Stanford), Steve Cappell (NIH), James Briscoe (The Francis Crick Institute), and members of the Teruel lab for helpful discussions and critical reading of the manuscript.

## SUPPLEMENTARY MATERIAL

- **Supplementary Video 1:** Dual-reporter OP9 cells induced to undergo adipogenesis by addition of the commonly-used DMI adipogenic stimulus. Daughter 1 (blue trace) and daughter 2 (red trace) only represent the product of the second mitosis (∼58 hr) in the time trace. The outline represents the outline of nuclear segmentation based on the H2B channel. APC/C reporter images and trace (gray) and H2B signal are shown only for daughter 1.
- **Supplementary Video 2:** Dual-reporter OP9 cells transfected also with a CDK2 live-cell sensor induced to undergo adipogenesis by addition of the commonly-used DMI adipogenic stimulus.
- **6 Supplementary Figures**

## SUPPLEMENTARY FIGURE LEGENDS

**Figure S1. Additional validation of citrine-PPARG as a marker of differentiation commitment**. (A) The PPARG derivative and integral values are poorer predictors of differentiation, Traces from Figure 1B were smoothed using a Butterworth filter and a five-point stencil was applied to the smoothed traces to estimate the PPARG derivative at each timepoint. The PPARG derivative traces were then analyzed in a similar manner as in Figure 1E. The bins range from −300 to 300 in intervals of 50.

(B) The PPARG integral at each time point was estimated using the trapz() function in MATLAB (MathWorks). The integral values were then analyzed in a similar manner as described in Figure 1E. The bins range from 0 to 5×105 in intervals of 104. Note that both the derivative and integral are poor predictors for the final differentiated state. In the case of the derivative, the range between the lowest predicted probability and the highest predicted probability is smaller than found in Figure 1E. The integral values, although a good predictor for a single time point, suffers from the lack of consistency across timepoints, and a single integral value cannot be used to separate undifferentiated and differentiated cells for all timepoints.

(C) The analysis in Figure 1E was done for more time points that span the duration of the experiments and presented as a heatmap where rows represent a timepoint and the columns represents the PPARG bins as described in Figure 1E. The dotted line represents the estimated PPARG threshold for the experiment. The expanded analysis shows that the PPARG threshold remains stable throughout the experiment.

(D) The switch from PPARG low to PPARG high occurs over a relatively short time window. Blue traces represent unaligned population medians of the PPARG abundance (top) and PPARG derivative (bottom). The red trace represents the PPARG abundance (top) and PPARG derivative (bottom) after computationally aligning all traces by the time when the PPARG threshold is crossed and is represented by the zero timepoint. The dashed lines highlight the time window around the peak in the PPARG derivative of the aligned traces and suggests that the switch between PPARG low and PPARG high states occurs over a short window of about 4 hours. All shaded regions represent the interquartile range (25th-75th percentiles).

(E) Unaligned correlations of endpoint markers of adipogenesis to PPARG over a typical 4-day DMI differentiation experiment. PPARG timecourses from the cells that differentiated after 96 hours in Figure 1D were averaged. At each time point, the Pearson correlation coefficient between the unaligned PPARG values, and the endpoint immunofluorescence values for adipocyte markers was calculated.

**Figure S2. The APC/C reporter behaves similarly to the Crl4-Cdt2-based sensor in marking the start of S-phase in OP9 cells**.

(A) Dual reporter cells infected with a Crl4-Cdt2 sensor tagged with iRFP670. Cells were stimulated to differentiate using the DMI cocktalk, and timecourses from individual cells are plotted to compare the dynamics of the APC/C reporter and Crl4-Cdt2 sensor.

(B) Comparison of median levels of the APC/C reporter and Crl4-Cdt2 sensors with t=0 marking the onset of S phase. Shaded regions represent the interquartile range (25th to 75th percentiles).

(C) Comparison of the median levels of APC/C reporter and Crl4-Cdt2 sensors at the onset of S-phases across multiple days of imaging. Shaded regions represent the interquartile range.

(A-C) In summary, the difference between the two probes in measuring the G1/S transition is only on the order of 1-2 hours at most, which would not change any of our conclusions in the context of our 4-day long experiments.

**Figure S3. Additional results supporting Figure 2**.

(A) Characterization of the plating conditions. In this study, we used subconfluent plating conditions (5K per well) instead of plating at confluent (15K per well) conditions that we normally use. Subconfluent plating increases the number of cell division events observed during adipogenesis and allows for easier cell tracking. The dual reporter cells were differentiated using the standard DMI cocktail. Left, The plot represents the fraction of cells that are considered past the PPARG threshold at each time point for both cell density conditions. Right, A comparison of the fraction of dual reporter cells in S/G2/M phases of the cell cycle, as assessed by the APC/C reporter, for both plating conditions.

(B) The number of mitotic events for both plating conditions are reported in the histogram as the fraction of cells observed with a given number of mitosis events. The 15,000 cells per well plating condition represents a standard differentiation protocol and yields high rates of differentiation and relatively low cell cycle activity. However, plating cells at a density of 5000 cells per well leads to a lower degree of differentiation and a higher degree of cell cycle activity. The live cell experiments in this manuscript are plated at a density of 5000 cells per well.

(C) Dilution through cell division does not significantly affect PPARG dynamics in differentiated cells. PPARG dynamics in differentiated cells were compared between cells that divided two (blue) or three (red) times in the span of the experiment, as indicated by the APC/C reporter (right). Additionally, the selected cells all completed the last mitosis at similar times. PPARG (left) and APC/C reporter (right) traces were computationally aligned to the last mitosis time. Bold traces represent median values and the shaded region represents the 95th confidence interval of the median.

(C) The trade-off between continued proliferation and differentiation exists even in cells that have been selected for undergoing exactly two divisions during the timespan of a 96-hour live-cell experiment.

(E) A CDK2 sensor (orange trace) was added to the PPARG and APC/C dual reporter cells to create triple reporter cells. Triple reporter cells were differentiated using the standard DMI protocol, and a representative trace is shown. The yellow dot represents the time when the cell reached the PPARG threshold and irreversibly committed to the differentiated state. Representative of 2 independent experiments.

(F) Scatter plot showing the CDK2 activity versus PPARG level in each single cell at every time point. The red dashed line represents the PPARG threshold.

**Figure S4. Significant cell-to-cell variability is apparent even when accounting for factors such as cell cycle phase when DMI was added, number of previous cell cycles, and refreshing the stimulus/serum**.

(A) Plot shows cells that differentiated in the experiment separated by what phase of the cell cycle the cells were in when DMI was added. Cells in which the adipogenic stimulus is added in G1 have fewer average additional divisions compared to cells where adipogenic stimuli are applied in S/G2/M.

(B) Analysis of the PPARG increase in cells with 0, 1 or 2 divisions before terminal cell differentiation.

(C) When the time of second mitosis is late, cells have on average slower increases in PPARG compared to cells where the second mitosis is earlier.

(D) Control experiment showing that replacement of DMI medium every 12 hours does not significantly change differentiation outcome.

**Figure S5. Validation of the siRNA knockdown efficiency when cells were transfected at 48 hours after induction of adipogenesis**.

(A) Cells were transfected with siRNA 48 hours after addition of the adipogenic DMI stimulus and knockdown efficiency was assessed 48 hours later (at the end of the 96-hour long time-lapse experiment). To validate the siRNA knockdown efficiency of p21 and CEBPA, cells were fixed with paraformaldehyde and immunostained for p21 or CEBPA levels.

(B) PPARG knockdown was assessed using the live cell citrine-PPARG signal.

**Figure S6. Adipogenic stimuli initiate a competition between proliferation and differentiation during a gradually extending G1 phase**.

(A) Histogram of the difference between G1_1_ and G1_2_ for each cell from Figure 5B.

(B) Plot of G1 duration versus how long a cell had been exposed to the adipogenic (DMI) stimulus at the start of G1 for each cell from (A). Red line marks average G1 duration of all cells.

(C) Analysis of timecourses in Figure 2B showing duration of G1 versus S/G2/M for differentiated and undifferentiated cells. Representative of 3 independent experiments.

(D-E) Analysis of timecourses of OP9 cells transfected with siRNA targeting cdc25c and cdc25c and induced to differentiate using a standard DMI protocol.

