## Supplemental Figures for "Molecular competition in G1 controls when cells simultaneously commit to terminally differentiate and exit the cell-cycle"

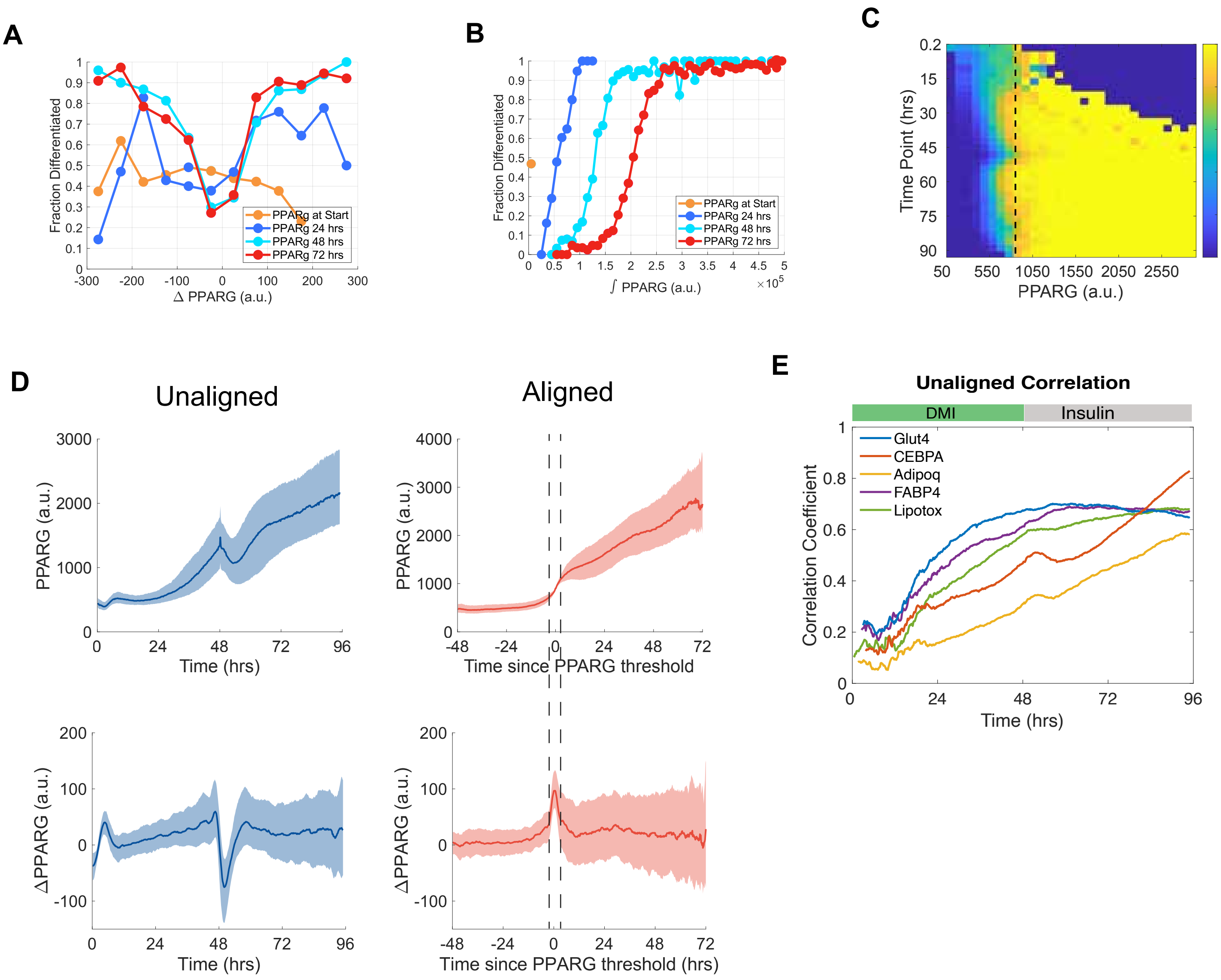


**Figure S1. Additional validation of citrine-PPARG as a marker of differentiation commitment.**

(A)The PPARG derivative and integral values are poorer predictors of differentiation, Traces from Figure 1B were smoothed using a Butterworth filter and a five-point stencil was applied to the smoothed traces to estimate the PPARG derivative at each timepoint. The PPARG derivative traces were then analyzed in a similar manner as in Figure 1E. The bins range from -300 to 300 in intervals of 50.

(B) The PPARG integral at each time point was estimated using the trapz() function in MATLAB (MathWorks). The integral values were then analyzed in a similar manner as described in Figure 1E. The bins range from 0 to 5x105 in intervals of 104. Note that both the derivative and integral are poor predictors for the final differentiated state. In the case of the derivative, the range between the lowest predicted probability and the highest predicted probability is smaller than found in Figure 1E. The integral values, although a good predictor for a single time point, suffers from the lack of consistency across timepoints, and a single integral value cannot be used to separate undifferentiated and differentiated cells for all timepoints.

(C) The analysis in Figure 1E was done for more time points that span the duration of the experiments and presented as a heatmap where rows represent a timepoint and the columns represents the PPARG bins as described in Figure 1E. The dotted line represents the estimated PPARG threshold for the experiment. The expanded analysis shows that the PPARG threshold remains stable throughout the experiment.

**(**D) The switch from PPARG low to PPARG high occurs over a relatively short time window. Blue traces represent unaligned population medians of the PPARG abundance (top) and PPARG derivative (bottom). The red trace represents the PPARG abundance (top) and PPARG derivative (bottom) after computationally aligning all traces by the time when the PPARG threshold is crossed and is represented by the zero timepoint. The dashed lines highlight the time window around the peak in the PPARG derivative of the aligned traces and suggests that the switch between PPARG low and PPARG high states occurs over a short window of about 4 hours. All shaded regions represent the interquartile range (25th-75th percentiles).

(E) Unaligned correlations of endpoint markers of adipogenesis to PPARG over a typical 4-day DMI differentiation experiment. PPARG timecourses from the cells that differentiated after 96 hours in Figure 1D were averaged. At each time point, the Pearson correlation coefficient between the unaligned PPARG values, and the endpoint immunofluorescence values for adipocyte markers was calculated.

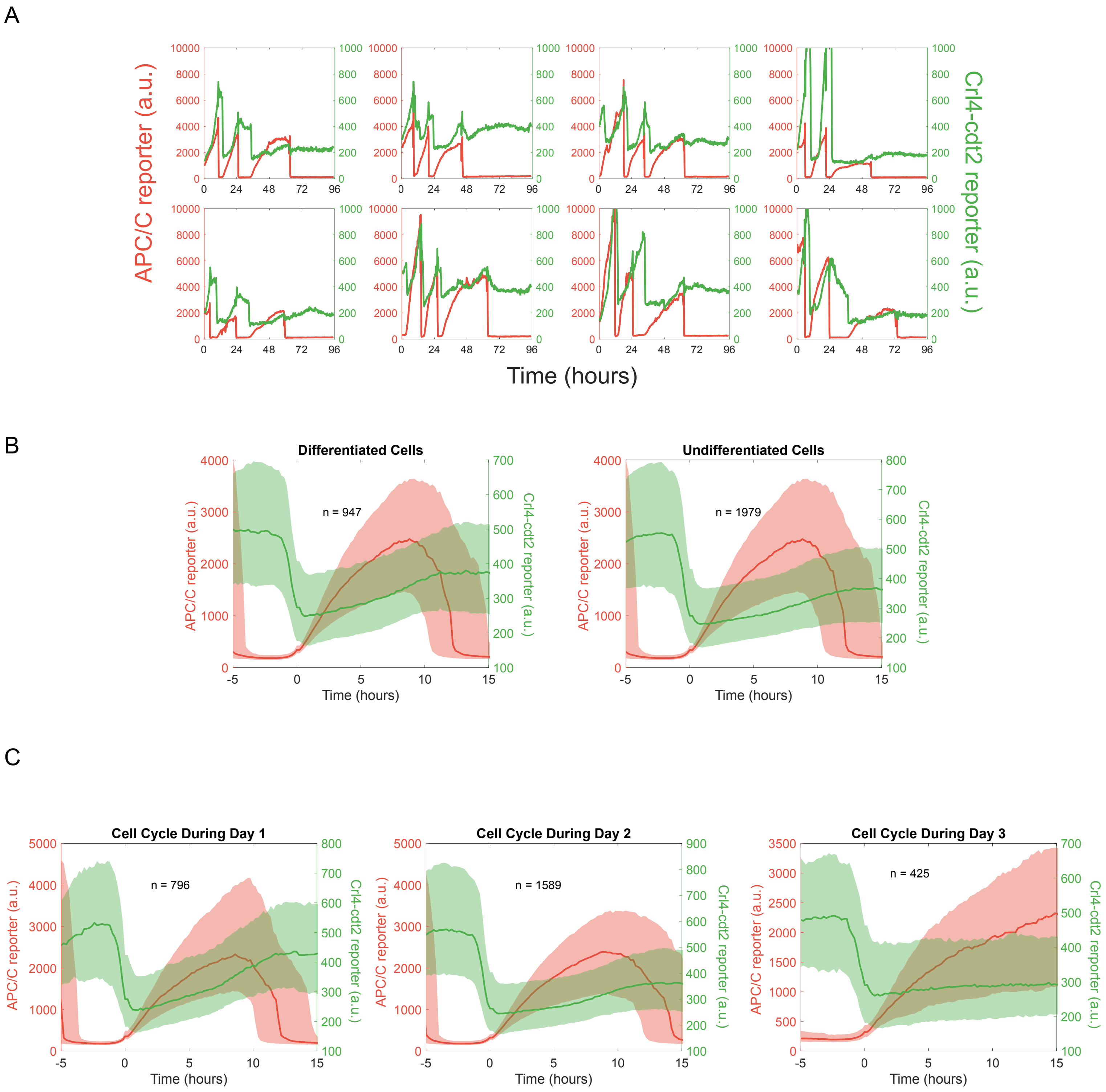


**Figure S2. The APC/C reporter behaves similarly to the Crl4-Cdt2-based sensor in marking the start of S-phase in OP9 cells.**
(A) Dual reporter cells infected with a Crl4-Cdt2 reporter tagged with iRFP670. Cells were stimulated to differentiate using the standard DMI 96-hour differentiation protocol, and timecourses from individual cells are plotted to compare the dynamics of the APC/C reporter and Crl4-Cdt2 sensor.
(B) Comparison of median levels of the APC/C and Crl4-Cdt2 reporter with t=0 marking the onset of S phase. Shaded regions represent the interquartile range (25th to 75th percentiles).
(C) Comparison of the median levels of APC/C and Crl4-Cdt2 reporters at the onset of S-phases across multiple days of imaging. Shaded regions represent the interquartile range.
(A-C) In summary, the difference between the two probes in measuring the G1/S transition is only on the order of 1-2 hours at most, which would not change any of our conclusions in the context of our 4-day long experiments.

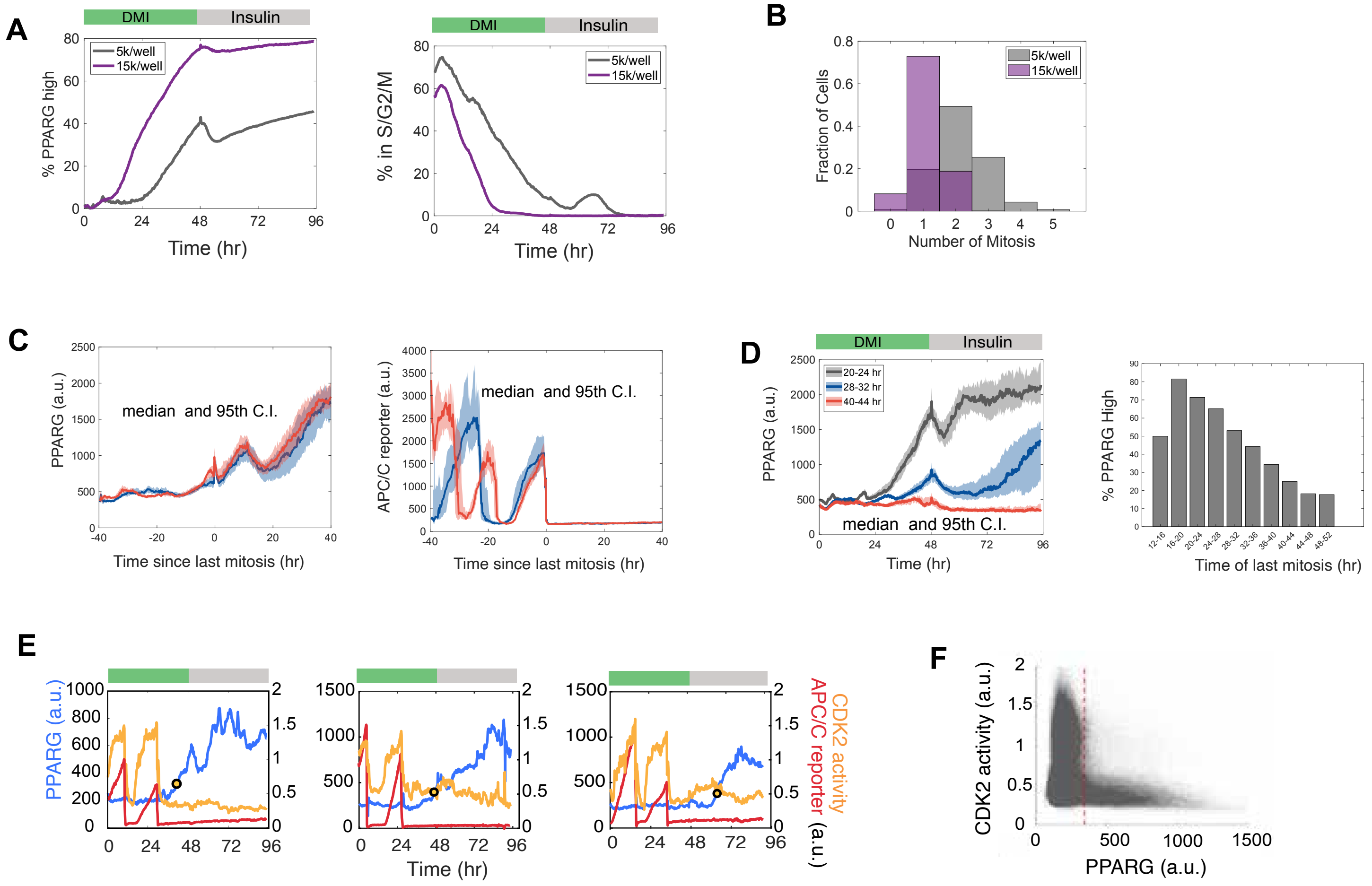


**Figure S3. Additional results supporting Figure 2.**

(A) Sub confluent plating conditions increase cell division events during adipogenesis. A comparison of the differentiation between dual reporter cells plated at two different cell densities. Dual reporter cells were differentiated using the standard DMI cocktail. *Left,* The plot represents the fraction of cells that are considered past the PPARG threshold at each time point for both cell density conditions. *Right,* A comparison of the fraction of dual reporter cells in S/G2/M phases of the cell cycle, as assessed by the APC/C reporter, for both plating conditions.

(B) The number of mitotic events for both plating conditions are reported in the histogram as the fraction of cells observed with a given number of mitosis events. The 15,000 cells per well plating condition represents a standard differentiation protocol and yields high rates of differentiation and relatively low cell cycle activity. However, plating cells at a density of 5000 cells per well leads to a lower degree of differentiation and a higher degree of cell cycle activity. The live cell experiments in this manuscript are plated at a density of 5000 cells per well.

(C) Dilution through cell division does not significantly affect PPARG dynamics in differentiated cells. PPARG dynamics in differentiated cells were compared between cells that divided two (blue) or three (red) times in the span of the experiment, as indicated by the APC/C reporter (right). Additionally, the selected cells all completed the last mitosis at similar times. PPARG (left) and APC/C reporter (right) traces were computationally aligned to the last mitosis time. Bold traces represent median values and the shaded region represents the 95th confidence interval of the median.

(D) The trade-off between continued proliferation and differentiation exists even in cells that have been selected for undergoing exactly two divisions during the timespan of a 96-hour live-cell experiment.

(E) A CDK2 sensor (orange trace) was added to the PPARG and APC/C dual reporter cells to create triple reporter cells. Triple reporter cells were differentiated using the standard DMI protocol, and a representative trace is shown. The yellow dot represents the time when the cell reached the PPARG threshold and irreversibly committed to the differentiated state. Representative of 2 independent experiments.

(F) Scatter plot showing the CDK2 activity versus PPARG level in each single cell at every time point. The red dashed line represents the PPARG threshold.

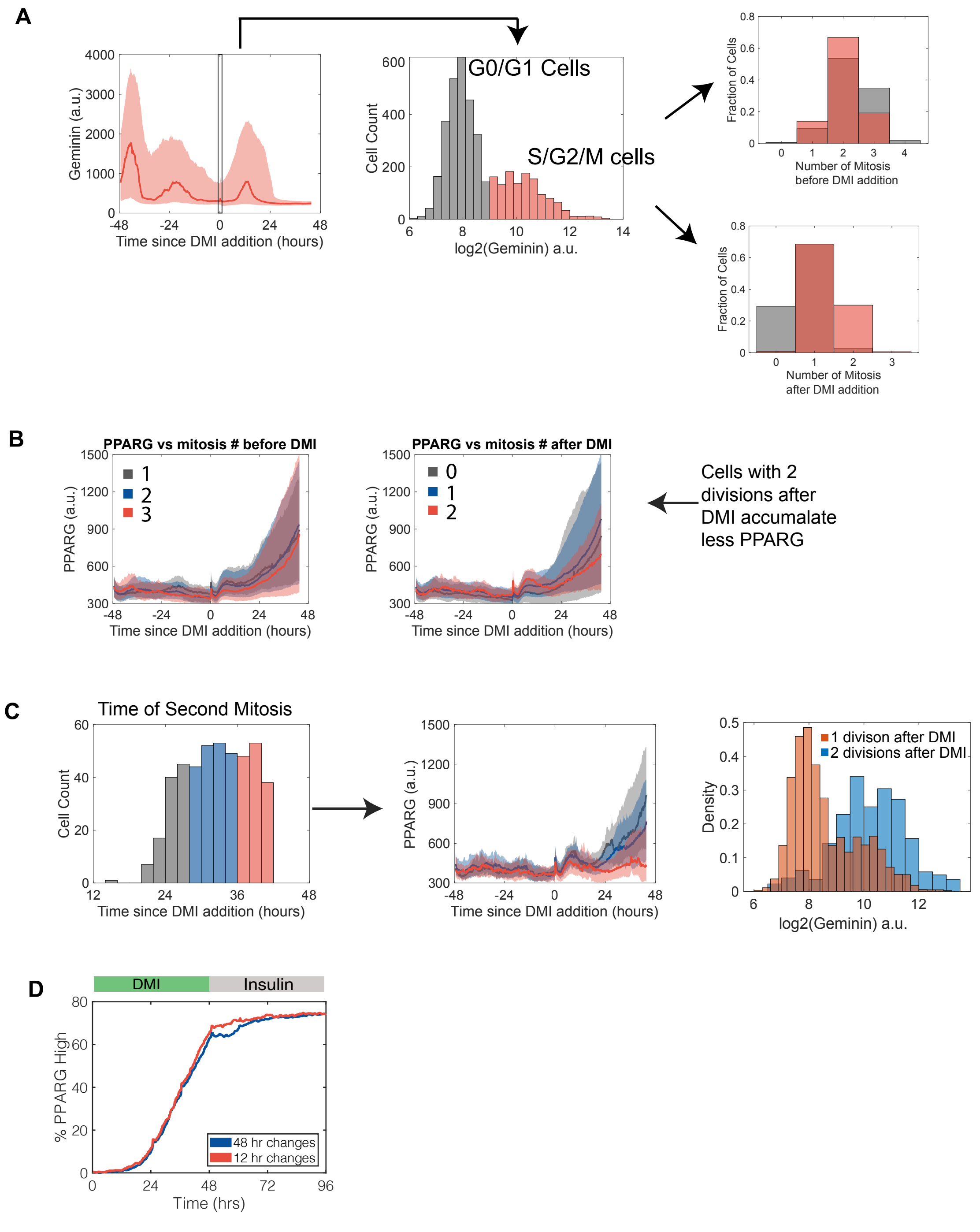


**Figure S4. Significant cell-to-cell variability is apparent even accounting for factors such as cell cycle phase when DMI was added, number of previous cell cycles, and refreshing the stimulus/serum.**
(A) Plot shows cells that differentiated in the experiment separated by what phase of the cell cycle the cells were in when DMI was added. Cells in which the adipogenic stimulus is added in G1 have fewer average additional divisions compared to cells where adipogenic stimuli are applied in S/G2/M.
(B) Analysis of the PPARG increase in cells with 0, 1 or 2 divisions before terminal cell differentiation.
(C) When the time of second mitosis is late, cells have on average slower increases in PPARG compared to cells where the second mitosis is earlier.
(D) Control experiment showing that replacement of DMI medium every 12 hours does not significantly change differentiation outcome.

A

B

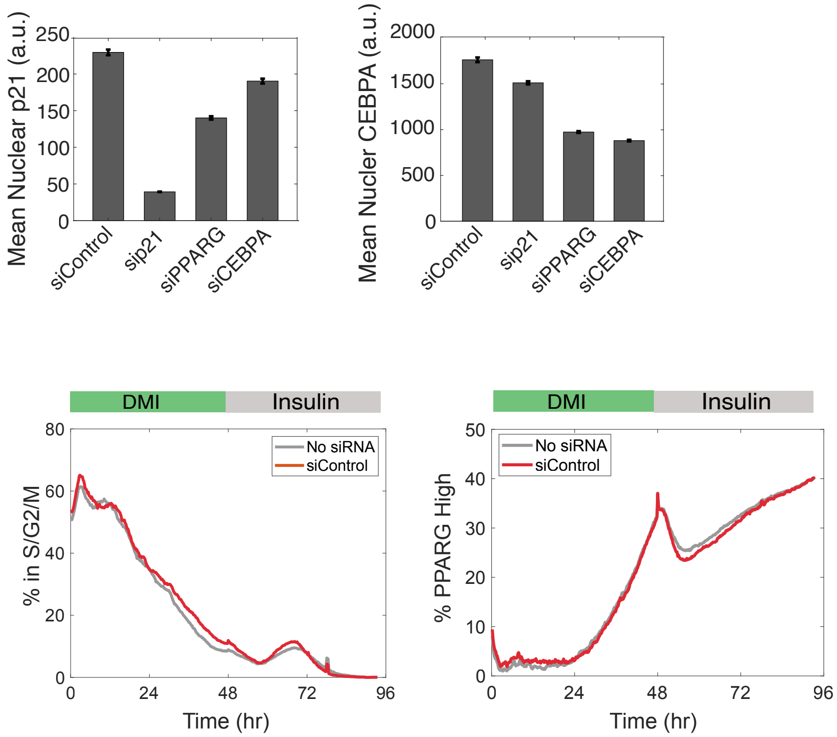


**Figure S5. Validation of the siRNA knockdown efficiency when cells were transfected at 48 hours after induction of adipogenesis.**

(A) Cells were transfected with siRNA 48 hours after addition of the adipogenic DMI stimulus and knockdown efficiency was assessed 48 hours later (at the end of the 96-hour long time-lapse experiment). To validate the siRNA knockdown efficiency of p21 and CEBPA, cells were fixed with paraformaldehyde and immunostained for p21 or CEBPA levels.

(B) PPARG knockdown was assessed using the live cell citrine-PPARG signal.

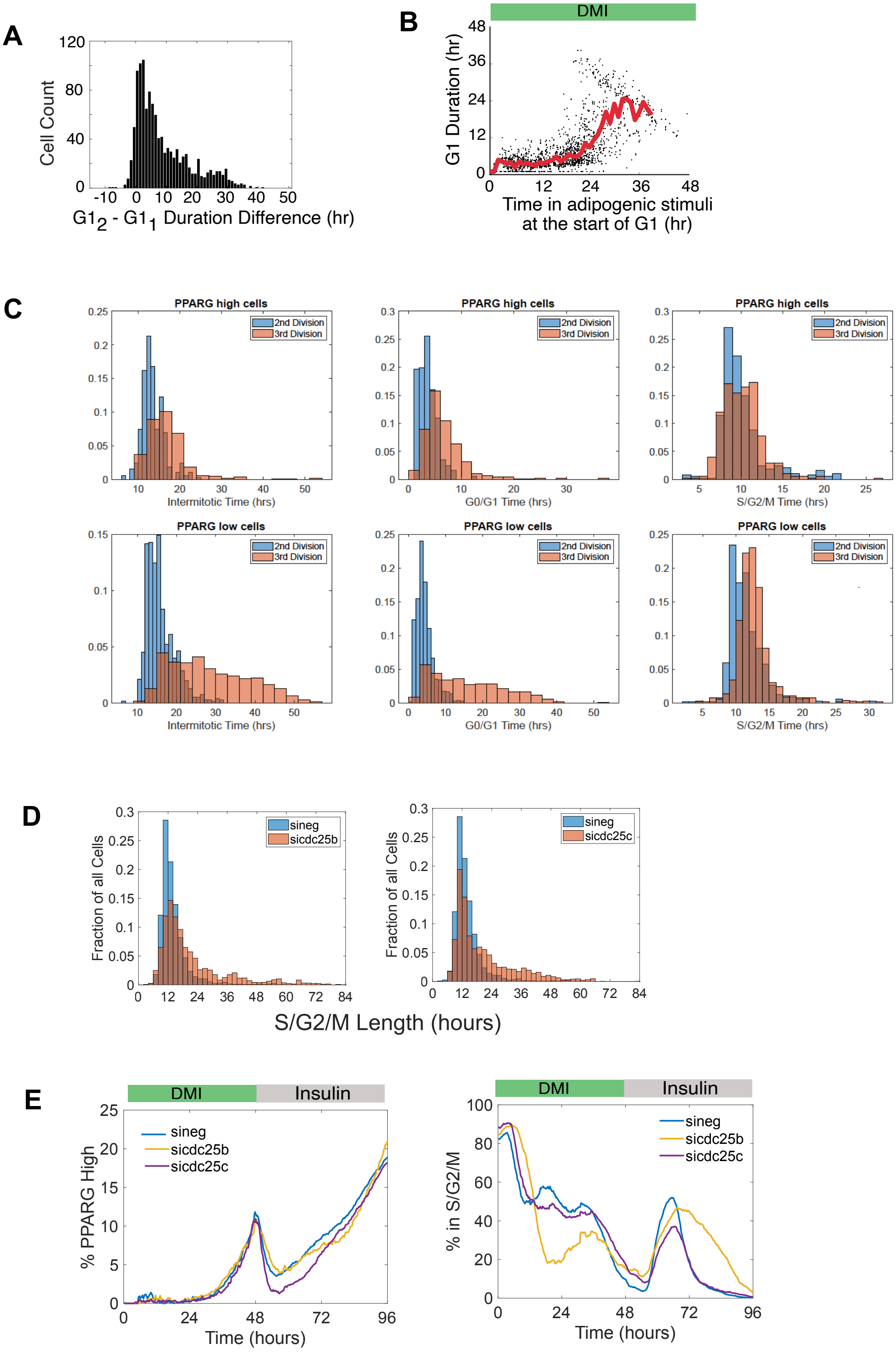


**Figure S6. Adipogenic stimuli initiate a competition between proliferation and differentiation during a gradually extending G1 phase.**
(A) Histogram of the difference between $G1_1$ and $G1_2$ for each cell from Figure 5B.
(B) Plot of G1 duration versus how long a cell had been exposed to the adipogenic (DMI) stimulus at the start of G1 for each cell from (A). Red line marks average G1 duration of all cells.
(C) Analysis of timecourses in Figure 2B showing duration of G1 versus S/G2/M for differentiated and undifferentiated cells. Representative of 3 independent experiments.
(D-E) Analysis of timecourses of OP9 cells transfected with siRNA targeting cdc25c and cdc25c and induced to differentiate using a standard DMI protocol.
